# Expansion of apical extracellular matrix underlies the morphogenesis of a recently evolved structure

**DOI:** 10.1101/686089

**Authors:** Sarah Jacquelyn Smith, Lance A. Davidson, Mark Rebeiz

## Abstract

One of the fundamental gaps in our knowledge of the evolution of novel structures is understanding how the morphogenetic processes that form these structures arise. Here, we traced the cellular development of a morphological novelty, the posterior lobe of *D. melanogaster*. We found that this genital outgrowth forms through an extreme increase in cell height. By examining the apical extracellular matrix (aECM), we uncovered a vast network associated with the developing genitalia of lobed and non-lobed species. We observed that cells which will form the posterior lobe show expanded expression of the aECM protein Dumpy which connects them to the ancestral aECM network. Further analysis demonstrated a required role for Dumpy in cell height increase during development. We propose that the aECM presents a rich reservoir for generating morphological novelty, in addition to highlighting a yet unseen role for aECM in regulating extreme cell height.

## Introduction

Biologists have long been mesmerized by the appearance of morphological novelties, new structures that appear to lack homologs in other species groups (Moczek, 2008; Günter et al., 2010). To understand the origins of these novel structures, significant effort has focused on determining how spatial and temporal patterning of genes are altered during evolution (Peter & Davidson, 2015; Rebeiz, et al., 2015; Wagner, 2014). This has indicated how developmental programs are often associated with morphological novelties, and they are frequently co-opted from other tissues. However, limited attention has been directed to how novel structures form at the cellular level. Understanding how a structure physically forms is important, as it can help explain which morphogenetic processes might be targeted during evolution. In addition, because most morphological novelties arose in the distant past, it is likely that the causative genetic changes will be obscured by additional changes scattered throughout relevant gene regulatory networks (Liu, Y., 2019). Hence, understanding the morphogenetic basis of a novelty is critical to identifying the most important aspects of the gene regulatory networks that contributed to its origin.

Most studies of morphogenetic evolution have focused on structures subject to diversification, illuminating processes that contributed to their modification, as opposed to origination. For example, studies of tooth morphogenesis have elucidated how both internal mechanisms, such as cell shape changes (Li et al., 2016), and external forces, such as the pressure from the surrounding jaw (Renvoisé et al., 2017) could be contributing factors in their diversification. An examination of the enlarged ovipositor of *Drosophila suzukii* revealed how a 60% increase in length was associated with increases in apical area and anisotropic cellular rearrangement (Green et al., 2019). In addition, differences in early morphogenetic mechanisms between distantly related species are observed in both the development of breathing tubes on the Drosophilid eggshell (Osterfield et al., 2015) and migration of sex comb precursors on *Drosophila* male forelegs (Atallah et al., 2009; Tanaka et al., 2009), together highlighting how rapid changes in morphogenetic mechanisms can evolve to form the same structure. Overall, these studies have illustrated how evolutionary comparative approaches can reveal morphogenetic processes critical to the sculpting of anatomical structures.

Morphogenesis is the product of both cell intrinsic processes, such as those conferred by the cytoskeleton or cell-cell junctions, and external forces from the environment in which the cell resides. Extracellular mechanics are relatively understudied compared to intracellular mechanics (Paluch & Heisenberg, 2009). An important component of the microenvironment of a cell is the extracellular matrix (ECM) which can be subdivided into two populations of ECM, the basal ECM and the apical ECM (aECM) (Brown, 2011; Daley & Yamada, 2013; Linde-Medina & Marcucio, 2018; Loganathan et al., 2016). While comparatively understudied, recent work has defined vital roles for aECM in the morphogenesis of structures, such as the *Drosophila* wing (Diaz-de-la-Loza et al., 2018; Etournay et al., 2015; Ray et al., 2015), denticles (Fernandes et al., 2010), and trachea (Dong et al., 2014), as well as in *C. elegans* neurons (Heiman et al., 2009; Low et al., 2019). Despite recent interest in the aECM, its role in the evolution of morphogenetic processes is currently unknown.

Genital traits represent a particularly advantageous system in which to study the morphogenetic basis of novel structures. The study of morphological novelty is often difficult because most structures of interest evolved in the distant past, rendering it difficult to understand the ancestral ground state from which the novelty emerged. Genitalia are noted for their rapid evolution (Eberhard, 1985), and thus bear traits among closely-related species that have recently evolved in the context of a tissue that is otherwise minimally altered. For example, the posterior lobe, a recently evolved anatomical structure present on the genitalia of male flies of the *melanogaster* clade (Kopp & True, 2002) (Figure 1A), is a three-dimensional outgrowth that is required for genital coupling (Frazee & Masly, 2015; Jagadeeshan & Singh, 2006; LeVasseur-Viens et al., 2015). Besides the posterior lobe, the genitalia of lobed and non-lobed species are quite similar in composition, providing an excellent context in which to examine the morphogenesis of the ancestral structures from which the posterior lobe emerged.

**Figure 1.**
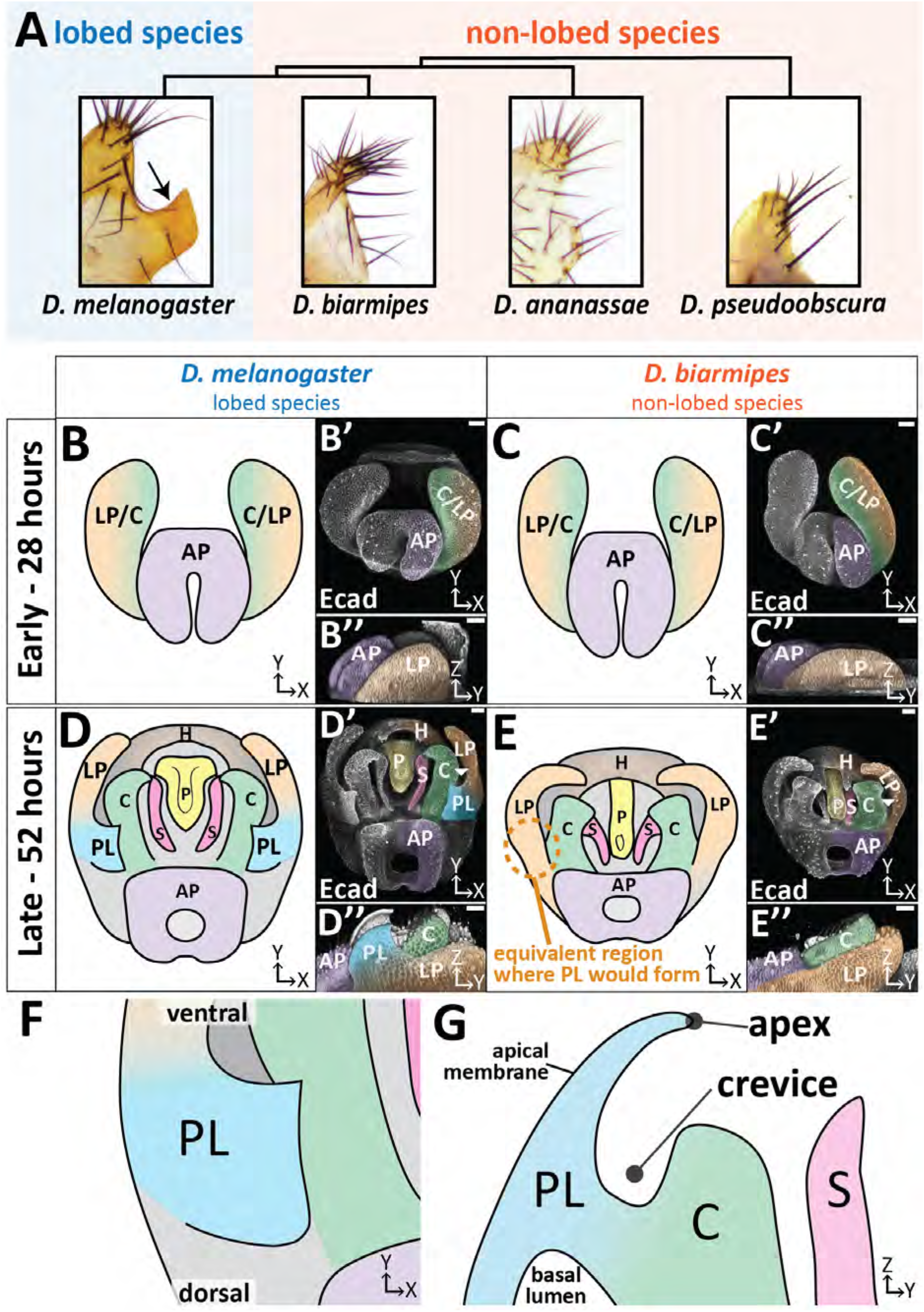
The posterior lobe a novel protrusion from the lateral plate of *D. melanogaster*. (A) Phylogenetic tree with representative bright-field images of adult cuticle of the lateral plate and posterior lobe (arrow). (B-E) Illustration, (B’-E’) maximum projection, and (B’’-E’’) three-dimensional projection of early (28 hours APF) and late (52 hours APF) developing genitalia showing the posterior lobe projecting form the lateral plate of *D. melanogaster* (D’’), but absent in *D. biarmipes* (E’’). Relevant structures are labeled: posterior lobe (PL), lateral plate (LP), clasper (C), sheath (S), phallus (P), anal plate (AP), and hypandrium (H). All max projections are oriented with ventral side towards to top and dorsal sides towards the bottom. (F) Zoomed in illustration of posterior lobe and (G) a cross-sectional/lateral view of the posterior lobe. The highest point of the lobe is the apex and the invagination between the lobe and the clasper is termed the crevice (G). Scale bar, 20μm.

Here, we find cell shape changes which increase cell height along the apico-basal axis drive morphogenesis of the posterior lobe. We investigated internal and external factors that might contribute to this height increase and find a correlation between the aECM protein Dumpy and the height of posterior lobe cells. Comparisons to non-lobed species uncovered the presence of a conserved aECM network on the genitalia that has expanded to cells that form the posterior lobe. Our work shows how the formation of a morphological novelty depends upon novel aECM attachments, integrating cells into a larger pre-existing aECM network.

## Results

### The posterior lobe grows from the lateral plate epithelium

The male genitalia of *Drosophila* is a bilaterally symmetrical anatomical structure which forms from the genital disc during pupal development. In adults, the posterior lobe protrudes from a structure called the lateral plate (also known as the epandrial ventral lobe (Rice et al., 2019)) (Figure 1A,D; Figure 1 - video 1). In *D. melanogaster*, prior to posterior lobe formation, the lateral plate is fully fused to a neighboring structure called the clasper (also known as the surstylus (Rice et al., 2019)) (Figure 1B) (Glassford et al., 2015). The lateral plate begins to separate from the clasper around 32 hours after pupal formation (APF) in *D. melanogaster* (Figure 1 – supplement 1). Approximately 4 hours later, the posterior lobe begins to project from the plane of the lateral plate and achieves its final shape by 52 hours APF (Figure 1D; Figure 1 - supplement 1). During posterior lobe development, cleavage of the lateral plate from the clasper continues, dropping the tip of the lateral plate behind the clasper and separating both tissues (Figure 1D; Figure 1 - supplement 1). Full separation of the lateral plate and clasper stops slightly above (ventral to) the posterior lobe (Figure 1 - supplement 1). By contrast, the lateral plate in the non-lobed species *D. biarmipes* remains flat throughout development, but all other morphogenetic events are very similar, forming on a schedule that is approximately 4 hours behind *D. melanogaster* (Figure 1C,E; Figure 1 - supplement 1).

### Posterior lobe cells increase in height to protrude from the lateral plate

To investigate which cellular behaviors are unique to lobed species, we examined how the posterior lobe grows from the lateral plate in both lobed and non-lobed species. First, we looked at cell proliferation, which commonly contributes to morphogenesis through patterned and/or oriented cell division (Heisenberg & Bellaïche, 2013), such as observed during branching morphogenesis in the lung where oriented cell division expands the bud before it bifurcates into two branches (Schnatwinkel & Niswander, 2013). During stages prior to the development of the posterior lobe morphogenesis, we observed widespread cell proliferation throughout the entire genital epithelium (Figure 2 - supplement 1). However, proliferation declines tissue-wide and all cell proliferation is essentially absent during posterior lobe development (Figure 2 - supplement 1). Similar dynamics in proliferation are also observed in non-lobed species (Figure 2 - supplement 1), suggesting that proliferation is not a major contributor to the morphogenesis of the posterior lobe.

**Figure 2.**
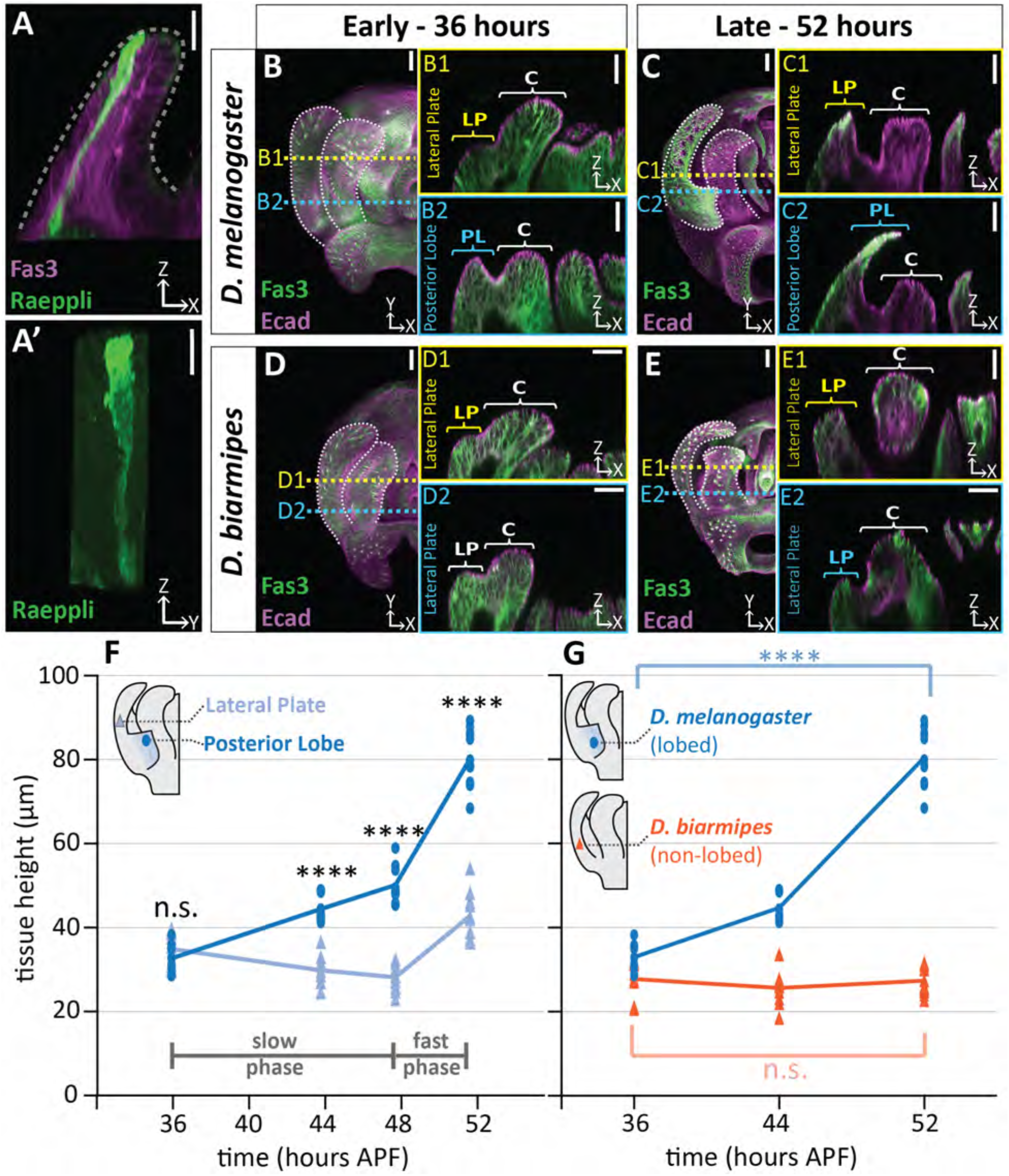
Posterior lobe cells increase in height to project out from the lateral plate. (A) A single cell in the posterior lobe labeled with Raeppli-mTFP1 (green) spans the height of the tissue labeled with lateral membrane marker fasciclin III (Fas3, magenta). Apical side of posterior lobe identified with dotted line. Sample is 44h after pupal formation (APF), but was heat shocked for 1 hour at 24h APF causing it to develop faster and more closely resembles a 48h APF sample. Scale bar, 10μm. n=4 (B-E) Maximum projections of early (36h APF) and late (52h APF) genital samples labeled with Fas3 (lateral membranes, green) and E-Cadherin (apical membranes, magenta). Location of respective cross sections indicated in yellow for lateral plate (B1-E1) and blue for posterior lobe (*D. melanogaster)* (B2-C2) or equivalent location in non-lobed species (*D. biarmipes*) (D2-E2). Scale bar, 20μm. (F) Quantification of tissue thickness of the lateral plate (light blue) and posterior lobe (dark blue). Illustration represents approximate location of cross-section that was used for tissue height measurement. Individual data points a presented; n=10 per each time point. (G) Quantification of tissue thickness of the posterior lobe in *D. melanogaster* (dark blue) and equivalent location in non-lobed species *D. biarmipes* (orange). Illustration represents approximate location of cross-section that was used for tissue thickness measurement. Individual data points presented; n≥9 per each time point. Statistical significance is indicated (unpaired t-test; ****p≤0.0001; n.s.=not significant p≥0.05). *D. melanogaster* tissue height measures in (G) are replotted from (F) to facilitate direct comparisons with *D. biarmipes*.

Next we tested the possibility that cell intercalation could contribute to posterior lobe morphogenesis. Such processes may play a role in tissue elongation (Guirao & Bellaïche, 2017; Tada & Heisenberg, 2012; Walck-Shannon & Hardin, 2014), such as in germ-band extension in *Drosophila* where directed cell intercalation results in a reduction in the number of cells on the anterior-posterior axis and an increase in the number of cells along the dorsal-ventral axis, elongating the tissue along the dorsal-ventral axis (Irvine & Wieschaus, 1994). To test this, we utilized live cell tracking during posterior lobe development. Initial observations of the outer face of the posterior lobe revealed few cell rearrangement events. When cell rearrangements did occur it was in response to a cell being removed from the apical surface (Figure 2 - video 1). Due to the limited number of cell rearrangement events observed during posterior lobe morphogenesis, cell intercalation does not appear to be a major driver of posterior lobe morphogenesis, causing us to instead examine changes in cell shape.

Changes to cell shape are quite common during tissue morphogenesis, as classically illustrated by the process of apical constriction that deforms tissues during many developmental processes (Lecuit & Lenne, 2007; Martin & Goldstein, 2014). To examine cell shape, we utilized the Raeppli system to label individual cells with a fluorescent marker (mTFP1) (Kanca et al., 2014). We observed that cells within the posterior lobe are tall and thin, spanning from the basal to the apical surface of the epithelium (Figure 2A). Because cells span the full thickness of this tissue, we can approximate the height of the tallest cells in the posterior lobe by measuring tissue thickness. For these measurements, we used the lateral plate as an in-sample comparison, since it represents the tissue from which the posterior lobe protrudes and should differ from the lobe in morphogenetic processes. We observed a pronounced increase in thickness of the posterior lobe compared to the lateral plate (Figure 2B-C,F; Figure 2 - supplement 2). The posterior lobe more than doubles in thickness with an average increase of 145.3% (+ 47.5μm), while the lateral plate only increases by 22.6% (+ 7.9μm) overall. In contrast, when non-lobed species are examined, no thickness changes are observed in the location where a posterior lobe would form, indicating that this increase in tissue thickness is unique to the posterior lobe (Figure 2B-E,G; Figure 2 - supplement 2). Interestingly, this increase in thickness is a dynamic process during development. During the first 12 hours of posterior lobe development the lateral plate thickness decreases by 5.1μm, but the posterior lobe increases in thickness by 16.5μm on average (Figure 2F). By contrast, during the last 4 hours of development, rapid increases in thickness occur in both the posterior lobe and lateral plate, which increase on average by 31.0μm and 14.6μm respectively (Figure 2F). These observations reveal a slow phase of cell height increase during the first 12 hours of posterior lobe development, and fast phase during the last four hours of posterior lobe development. Together this data suggests that the cells of the posterior lobe undergo an extreme cell shape change to increase in length along their apico-basal axis, driving the posterior lobe cells to project out of the plane of the lateral plate.

### Cytoskeletal components increase in concentration in posterior lobe cells

Elongation of cells along their apico-basal axes appears to be a major contributor to posterior lobe formation. To understand potential internal forces contributing to this cell shape change, we examined the organization of cytoskeletal components. As expected for a polarized epithelium, we found F-actin strongly localized to the apical cortex overlapping with E-cadherin throughout the entire genitalia (Figure 3A). In contrast with the adjacent tissues, F-actin is also concentrated along the apico-basal axis of posterior lobe cells (Figure 3A). This F-actin localization was unique to the posterior lobe, as it is less intense in neighboring structures, such as the lateral plate, clasper, and sheath, as well as in non-lobed species (Figure 3A; Figure 3 - supplement 1). Next we evaluated microtubules by examining two post-translational modifications that appear on tubulin, acetylation of α-tubulin on lysine40, a stabilizing modification (Roll-Mecak, 2019; Xu et al., 2017), and tyrosinated tubulin, which has been associated with rapid microtubule turnover (Roll-Mecak, 2019; Webster, Gundersen et al., 1987). In the posterior lobe, acetylated tubulin levels are highest at the apex of the posterior lobe and weaken towards the basal side of the lobe (Figure 3B-C). Compared to other structures in the genitalia, acetylated tubulin is greatly increased specifically in the posterior lobe (Figure 3B-C). In contrast, the levels of acetylated tubulin in non-lobed species are similar throughout the genitalia (Figure 3 - supplement 1). We found tyrosinated tubulin has a more consistent signal along the entire apico-basal axis in the posterior lobe (Figure 3B&D). The amount of tyrosinated tubulin in posterior lobe cells is increased compared to neighboring structures, but is weaker relative to the observed differences in acetylated tubulin. In non-lobed species the levels of tyrosinated tubulin are consistent across the entire genitalia (Figure 3 - supplement 1). Collectively, these results suggest that changes in assembly and/or dynamics of both F-actin and microtubule cytoskeletal networks could be contributing factors in changing the shape of posterior lobe cells to increase its height along the apico-basal axis.

**Figure 3.**
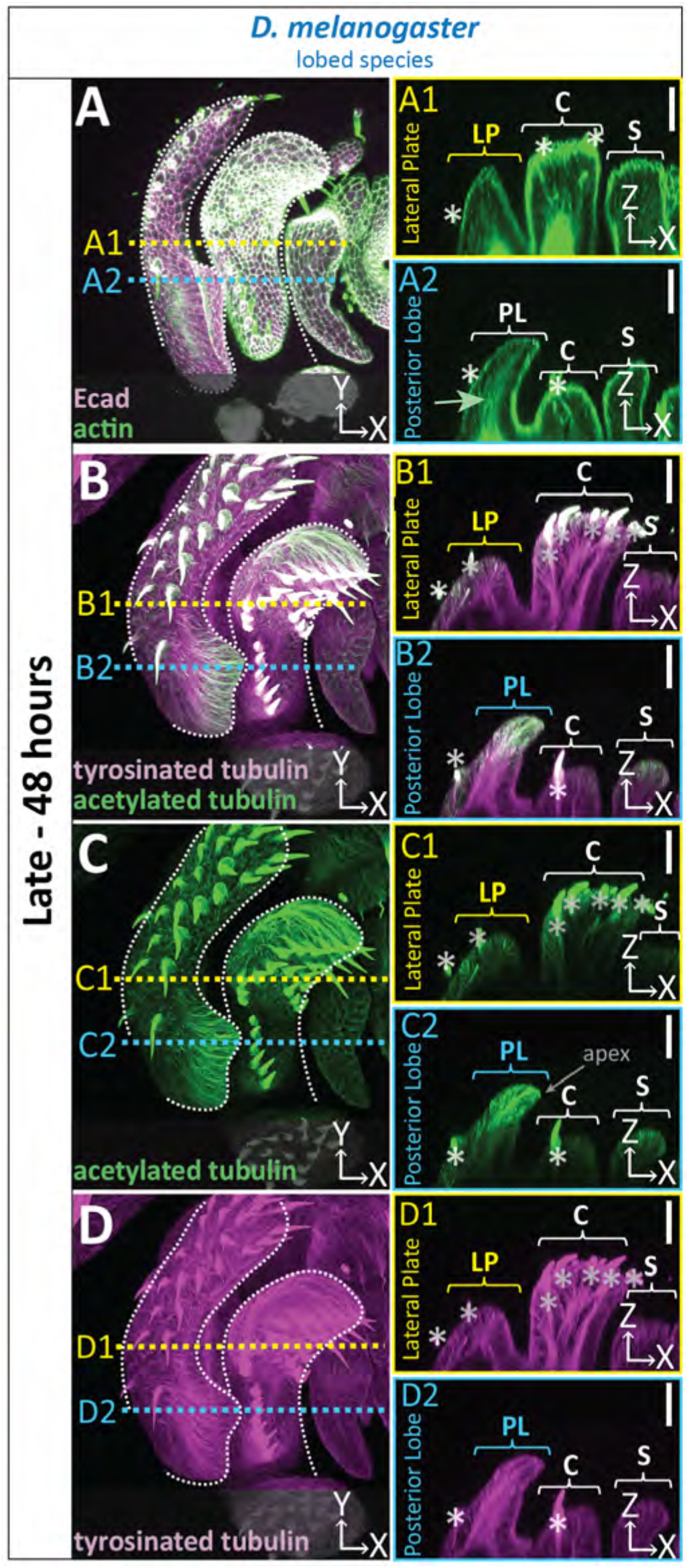
Cytoskeletal components are concentrated in posterior lobe cells. (A-D) Maximum projection, and respective cross-sections of late (48h APF) genital samples of the lobed species *D. melanogaster* labeled with F-actin/phalloidin and Ecad (A), acetylated tubulin (B,C), and tyrosinated tubulin (B,D). Location of respective cross sections indicated in yellow for lateral plate (A1-D1) and blue for posterior lobe (A2-D2). Cross-sections are maximum projections of a restricted 5.434μm thick section to provide a complete view of cytoskeletal components along the apico-basal axis. All cross-sections are oriented with apical side at the top and basal side at the bottom. Asterisk identifies bristles which have high levels of F-actin and tubulin. Bright basal signal in A1 and A2 are fat bodies. Bottom layers were removed in panel A to remove fat body signal which overwhelmed other details. (B-D2) Panels C and D show separate channels of panel B. Relevant structures labeled: Posterior lobe (PL), lateral plate (LP), clasper (C), and sheath (S). Scale bar, 20μm. n≥ 3 per experiment.

### An apical extracellular matrix associates with posterior lobe cells

In addition to investigating cell autonomous mechanisms leading to increases in tissue thickness, we also sought to identify sources of external forces which could play a role in posterior lobe morphogenesis. Extrinsic roles for the basal and apical extracellular matrix have been established in the pupal wing of D. melanogaster (Diaz-de-la-Loza et al., 2018; Etournay et al., 2015; Ray et al., 2015). We first attempted to characterize the basal ECM by analyzing a GFP-tagged version of Collagen IV (Viking:GFP). We observed that Viking:GFP, while present at very early stages of genital morphogenesis, is weakly present during posterior lobe formation across the entire genitalia (Figure 4 - supplement 1), suggesting that minimal basal ECM is present at this time point. To further test for the presence of basal ECM, we examined another basal ECM component, Perlecan (Perlecan:GFP), and also observed weak signal (Figure 4 – supplement 1). Together, this data suggests that the basal ECM is globally decreased in the genitalia during early pupal development, such that it is very weak during posterior lobe morphogenesis.

**Figure 4.**
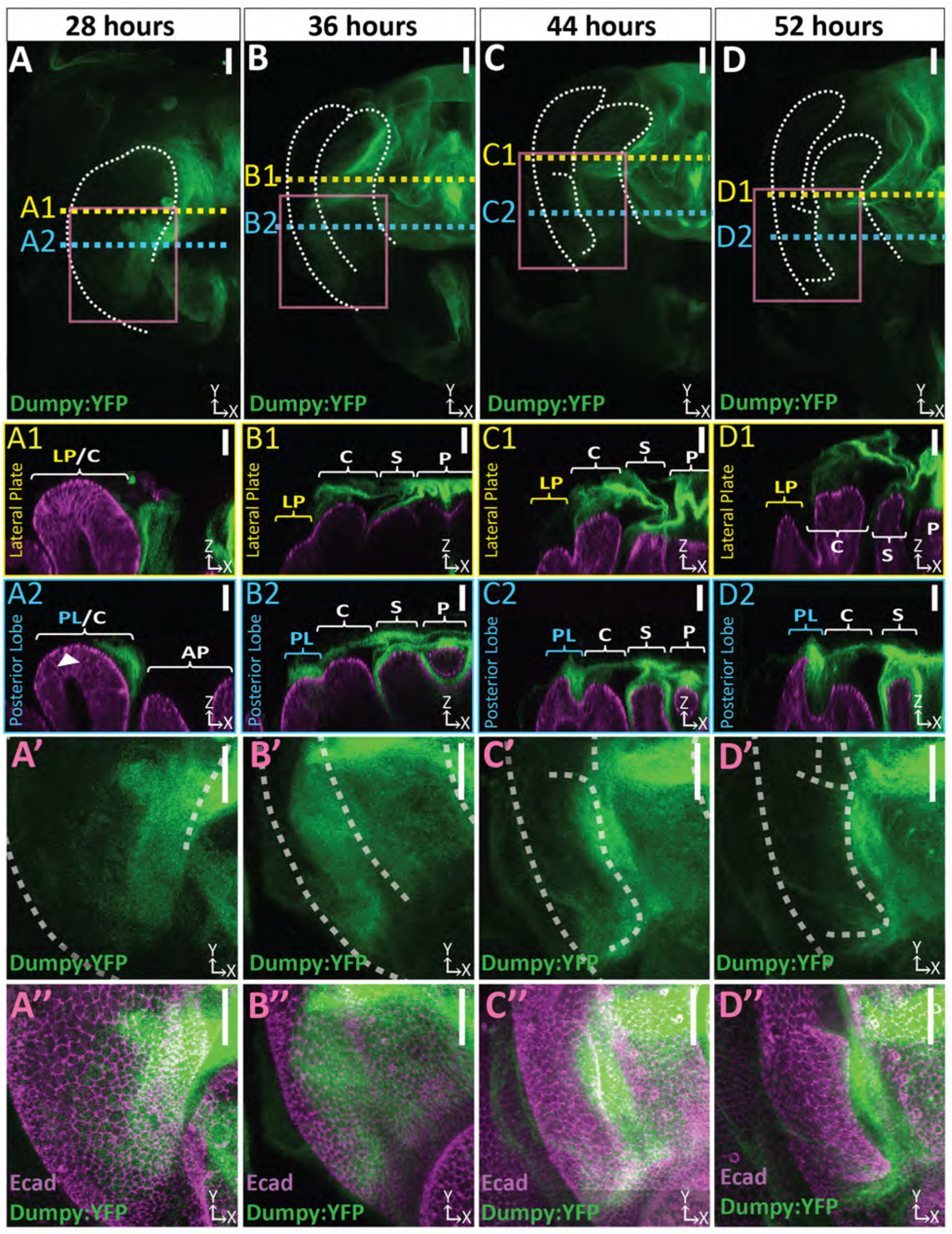
Dumpy deposition is correlated with posterior lobe development. (A-D) Maximum projection and (A’-B’’) respective zoom, indicated with pink box, labeled with Dumpy:YFP (green) and Ecad (magenta) for each time point. Location of respective cross sections indicated in yellow for lateral plate (A1-D1) and blue for posterior lobe (A2-D2). Arrowhead in (A2) indicates future posterior lobe cells. Cross-sections are oriented with apical side at the top and basal side at the bottom. Relevant structures labeled: Posterior lobe (PL), lateral plate (LP), clasper (C), sheath (S), and phallus (P). Scale bar, 20μm. n≥ 4 per experiment. Images were independently brightened to show relevant structures.

We next sought to determine if an aECM is present, and if so, whether it could potentially influence posterior lobe morphogenesis. A major component of the aECM is Dumpy, which is a gigantic (2.5 MDa) zona pellucida domain-containing glycoprotein (Wilkin et al., 2000). We examined a line in which Dumpy is endogenously tagged with a Yellow Fluorescent Protein (Dumpy:YFP). Dumpy:YFP forms a complex three-dimensional network over the pupal genitalia and is closely associated with cells of the posterior lobe (Figure 4; Figure 4 - video 1). At certain points in the genitalia, this aECM network of Dumpy can extend up to 39.4 μm on average above the cells, which is taller than the thickness of posterior lobe cells at the beginning of development (Figure 4 - supplement 2). The intricate complex morphology of this aECM network is hard to fully appreciate in flattened images due to its three-dimensional shape and spatially varying levels of Dumpy:YFP, making it difficult to see weaker populations of Dumpy without over-saturating more concentrated deposits.

In late pupal wing development, Dumpy anchors the wing to the surrounding cuticle, preventing the tissue from retracting away from the cuticle, which is important to properly shape the wing (Etournay et al., 2015; Ray et al., 2015). This same mechanism has been hypothesized to also occur in the leg and antennae (Ray et al., 2015), however, in the posterior lobe we do not find discrete anchorage points to the cuticle. Instead, we observed a large tether of Dumpy emanating from the anal plate and connecting with the pupal cuticle membrane that encases the entire pupa (Figure 4 - supplement 3, video 2) (Bainbridge & Bownes, 1981). This tether does not come in direct contact with posterior lobe associated Dumpy or other nearby structures such as the lateral plate, clasper, sheath, or phallus, suggesting that if Dumpy is contributing to posterior lobe evolution and morphogenesis, it is likely through a mechanism which does not depend on a direct mechanical linkage with the overlying pupal cuticle.

To investigate the role that Dumpy may play in posterior lobe morphogenesis, we examined its localization throughout development. Prior to posterior lobe development, future cells of the lobe lack apical Dumpy, and yet an intricate network associated with the clasper is observed (Figure 4A). However, from the early stages of posterior lobe development, as it first protrudes from the lateral plate, we observe large deposits of Dumpy associated with future lobe cells (Figure 4B). These deposits persist throughout most of its development (Figure 4C), becoming more restricted to the apex of the posterior lobe towards the end of posterior lobe development (Figure 4D). Throughout development, the posterior lobe associated Dumpy population is connected to the complex network of Dumpy attached to more medial structures such as the phallus (Figure 4 A2-D2), indicating that the posterior lobe is interconnected via the aECM with nearby structures (Figure 4). In contrast to the posterior lobe, the lateral plate has minimal Dumpy associated with it (Fig. 4 A1-D1). Only when we oversaturate the Dumpy:YFP signal can we observe a weak population of Dumpy associated with the lateral plate (Figure 4 - supplement 4). Together, this indicates that the cells of the posterior lobe and the lateral plate substantially differ in the levels of associated Dumpy, suggesting a potential role in the morphogenesis of the posterior lobe.

### Expansion of Dumpy expression is correlated with the evolution of the posterior lobe

The association of the posterior lobe with Dumpy suggests that changes in the expression of *dumpy* may have been significant during the evolution of the posterior lobe. To test if posterior lobe-associated Dumpy is a unique feature of species which produce a posterior lobe, we compared the spatial distribution of its mRNA in *D. melanogaster* with *D. biarmipes*, a species which lacks this structure. Early in pupal genital development at 32 hours APF we observe very similar expression patterns of *dumpy* between *D. melanogaster* and *D. biarmipes*, with expression at the base of the presumptive lateral plate-clasper (Figure 5A, Figure 5 - supplement 1). From 36 to 40 hours APF, when the posterior lobe begins to develop, this pattern becomes restricted to a small region at the base of the lateral plate and clasper, near the anal plate in *D. biarmipes*, but is expanded in lobed species (Figure 5B, Figure 5 - supplement 1). By 44 hours APF, expression of *dumpy* is reduced in the posterior lobe, as well as in non-lobed species, with strongest expression associated with the clasper in *D. biarmipes* (Figure 5A-B, Figure 5 - supplement 1). Overall, these results indicate that expression of *dumpy* is expanded in a lobed species and correlates with the timing of the posterior lobe’s formation. In addition, considering that the developmental timing of *D. biarmipes* lags behind *D. melanogaster* by approximately 4 hours (Figure 1 - supplement 1), this suggests that *dumpy* expression becomes restricted during an earlier developmental period in the non-lobed species *D. biarmipes*.

**Figure 5.**
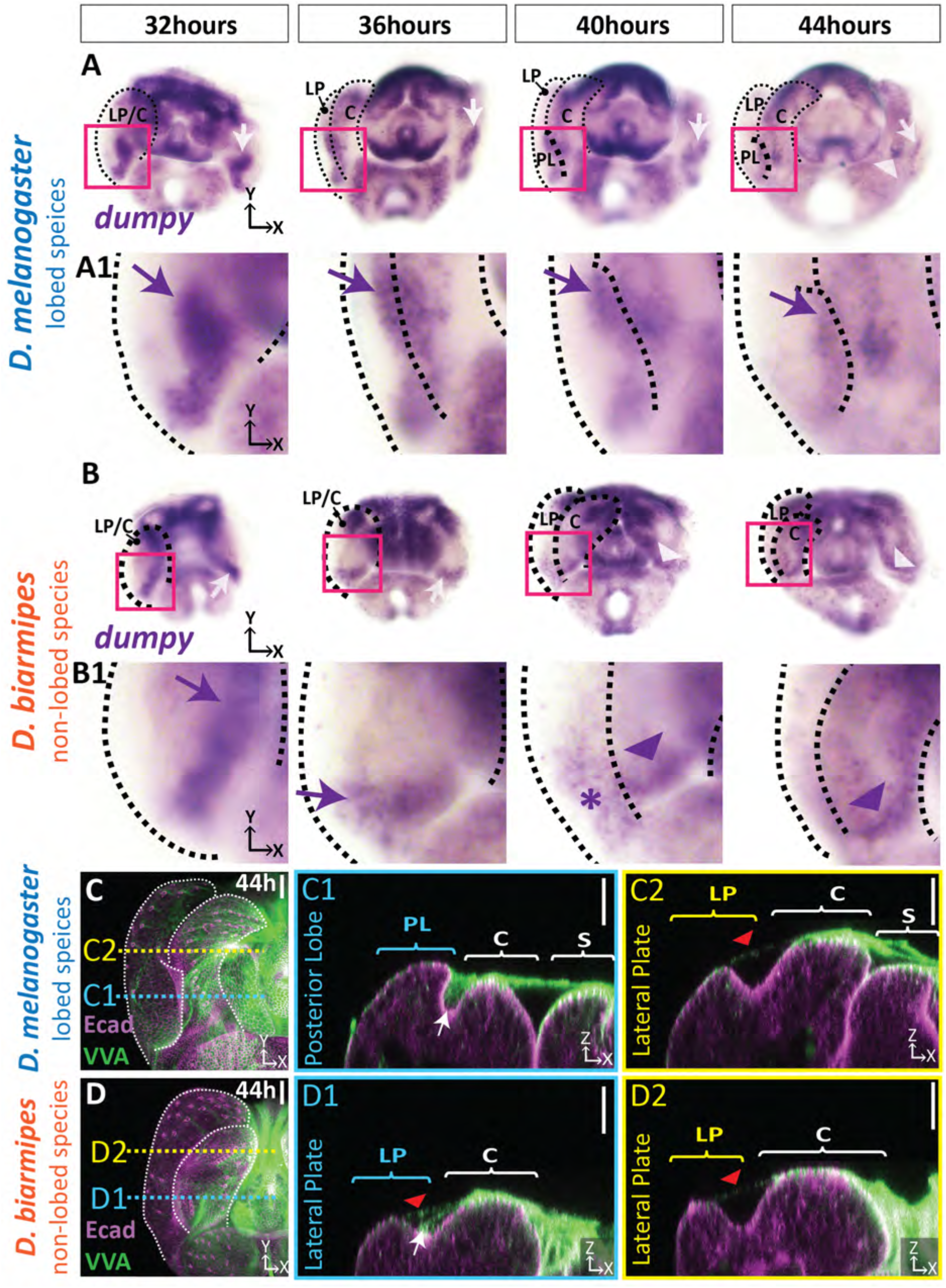
aECM is spatially expanded in lobed species compared to non-lobed species. (A-B) *in situ* hybridization for *dumpy* mRNA in the lobed species *D. melanogaster* (A) and the non-lobed species *D. biarmipes* (B). Pink box outlines location of zoom in for A1 and B1. Posterior lobe associated expression highlighted with arrow (purple/white) for strong expression, asterisk for weak expression, and arrowhead for clasper-specific expression. Expression observed in *D. melanogaster* at 44 hours APF is not present in all samples (see Figure 5 – supplement 1). (C-D) aECM is labeled with Vicia villosa lectin (VVA; green) and apical membrane labeled with Ecad (magenta) at 44 hours APF in *D. melanogaster* (C) and *D. biarmipes* (D). Location of respective cross sections indicated in yellow for lateral plate (C2-D2) and blue for posterior lobe in *D. melanogaster* (C1) and corresponding position in *D. biarmipes* (D1). All cross-sections are oriented with apical side at the top and basal side at the bottom. White arrows highlight the crevice localization between the lateral plate and clasper, which the aECM fills in *D. melanogaster* (C1), but only a weakly stained strand-like structure of aECM appears in *D. biarmipes* (D1). Tendrils of aECM can also be observed connecting to the lateral plate in both species (red arrowheads). Relevant structures labeled: Posterior lobe (PL), lateral plate (LP), clasper (C), sheath (S), and phallus (P). Scale bar, 20μm. n=at least 5 per experiment.

Although, it appears that the expression of *dumpy* has expanded in *D. melanogaster*, Dumpy is an extracellular protein, and cells expressing its mRNA may not correlate with its ultimate protein abundance or localization. Since an antibody for Dumpy is not available, we adapted lectin staining protocols which can detect glycosylated proteins like Dumpy in order to compare the distribution of aECM in species which lack posterior lobes. We found that fluorescein conjugated *Vicia villosa* lectin (VVA), which labels *N*-acetylgalactosamine (Tian & Ten Hagen, 2007), approximately recapitulated Dumpy:YFP in *D. melanogaster.* VVA strongly associates with the posterior lobe, shows trace association with the lateral plate, and roughly mirrors the complex three-dimensional shape of the Dumpy aECM network covering the center of the genitalia (Figure 5C). When we examined VVA in the non-lobed species *D. biarmipes*, we observed strong VVA signal over the center of the genitalia with weak connections to the tip of the lateral plate, similar to what we observe in *D. melanogaster* (Figure 5 C-D). In contrast, we only found a weak strand-like structure emanating from the clasper and connecting to the crevice between the lateral plate and clasper where the presumptive posterior lobe would form (Figure 5D). These results correlate with our *in situ* results, where we observe high expression at the center of the genitalia and weak expression of *dumpy* at the base between the clasper and lateral plate in *D. biarmipes*, which may be responsible for forming the weak aECM connection from the clasper to the crevice. Further, we observed similar staining patterns in an additional non-lobed species, *D. ananassae* (Figure 5 - supplement 2). Collectively, these data suggest that an ancestral aECM network was associated with the central genital structures, including the phallus, sheath, and clasper, and a weak association in the crevice next to prospective posterior lobe cells. During the course of evolution, expression of *dumpy* has expanded to integrate cells of the posterior lobe, creating a prominent connection to the aECM network.

### Dumpy is required for proper posterior lobe formation

Thus far, we observed a strong association of the aECM with cells that form the posterior lobe, a trait which is much less pronounced in non-lobed species. To determine if Dumpy plays a role in posterior lobe formation, we next employed transgenic RNAi to knock down its expression. Previous studies of *dumpy* characterized a VDRC RNAi line that is effective at reducing its function (Ray et al., 2015). We used a driver from the *Pox neuro* gene (Boll & Noll, 2002) to reduce *dumpy* levels in the posterior lobe. This resulted in a drastic decrease in the size and alterations to the shape of the posterior lobe compared to a control RNAi (Figure 6). In *dumpy* knockdown individuals, we observe a variable phenotype, and even within single individuals, the severity of phenotype differs between left and right posterior lobes (Figure 6A; Figure 6 - supplement 1). Knockdown was completed at both 25°C and 29°C, as higher temperatures increase the efficacy of the Gal4/UAS system (Duffy, 2002). At higher temperatures, the *dumpy* knockdown phenotype trended towards more severe defects (Figure 6B). Together, these results suggest that posterior lobe development is sensitive to levels of *dumpy*, and that *dumpy* plays a vital role in shaping the posterior lobe.

**Figure 6.**
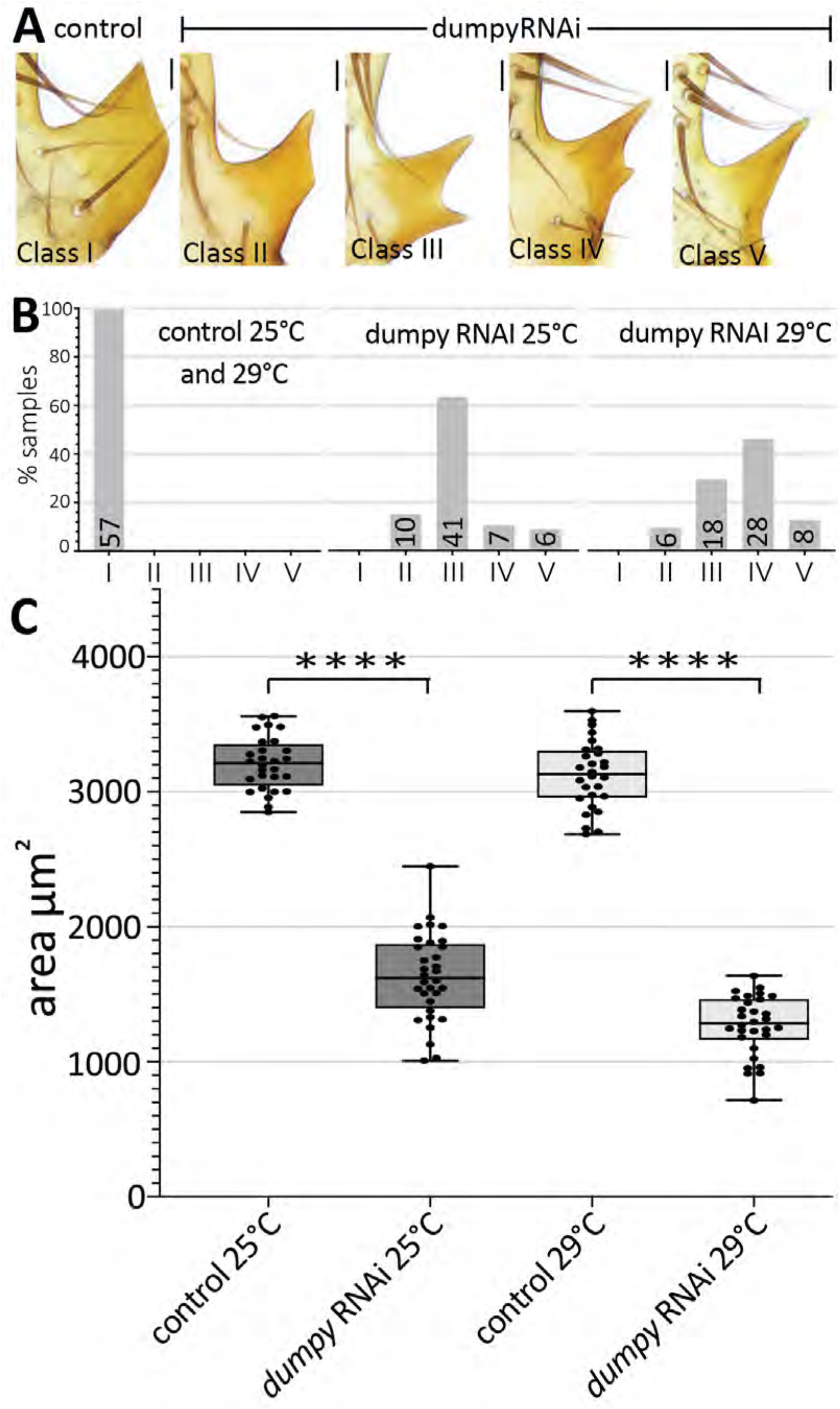
Dumpy is required for proper posterior lobe shape. (A) Range of adult posterior lobe phenotypes produced by control (*mCherry* RNAi) and *dumpy* RNAi animals. Phenotypic classes defined from wild type (I) to most severe (V). Scale bar, 20μm. (B) Percentage of posterior lobes in each class for control, *dumpy* RNAi at 25°C, and *dumpy* RNAi at 29°C. (C) Quantification of area of adult posterior lobes of *mCherry* RNAi (control) and *dumpy* RNAi at 25°C and 29°C. Statistical significance between each temperature indicated (unpaired t-test; ****p≤0.0001).

### Correlation of Dumpy deposition and cell height in the posterior lobe

We next sought to determine when during development *dumpy* knockdown influences the morphogenetic progression of the posterior lobe. This was important because we observed both a slow and a fast phase of lobe development (Figure 1F), and also reasoned that posterior lobe cells secrete cuticle once they have adopted their final adult conformations, of which any of these phases could represent a critical Dumpy-dependent stage of development. We found that *dumpy* knockdown individuals manifest phenotypes very early on (Figure 7A) and continue to show abnormal lobe development through the end of its formation (Figure 7B). Interestingly, differences in the height of cells on the ventral side of the posterior lobe are not observed between control and *dumpy* knockdown treatments, instead defects in cell height are observed in the more dorsally-localized cells of the posterior lobe (Figure 7A-B). This correlates with the phenotypes of the adults in the *dumpy* knockdown in which the ventral tip is usually of normal height with defects observed towards the dorsal side (Figure 6A). However, this phenotype appears counterintuitive, as Dumpy protein normally associates along the entire posterior lobe, so why does the tip of the posterior lobe develop to normal height when Dumpy is absent? To better understand this phenotype, we examined Dumpy:YFP localization in the *dumpy* knockdown background. We observed weak association of Dumpy with the tallest cells on the ventral side of the posterior lobe both in early (Figure 7D n=5/5 samples) and late (Figure 7F n=4/5 samples) stages compared to control animals. In contrast, no Dumpy was observed in contact with the short cells on the dorsal side (Figure 7D & F). Together, this highlights a correlation between the height of posterior lobe cells and presence of dumpy. One of our late samples lacks a Dumpy connection to the ventral cells, correlating with our observation that not all adult samples are fully extended on the ventral side (Figure 7 - supplement 1). This suggests that ventral cell connections to the Dumpy aECM network may be lost late in development, ultimately causing a shortening of these cells. In addition, we observed more severe phenotypes of *dumpy* knockdown in the *dumpy-yfp background* (compared to the *dumpy*-WT background alone), suggesting that Dumpy:YFP is a mild hypomorph (not shown). We also observed at early time points highly variable strands of Dumpy in the middle of the lobe (between the ventral and dorsal sides) (Figure 7 - supplement 2). These strands visually resembled the weak strands of VVA observed in *D. biarmipes* (Figure 5D), in that they emanate from the clasper and connect to the crevice between the posterior lobe and clasper. Overall, the most pronounced phenotypic defects manifest in regions with the strongest reduction in Dumpy aECM deposition, implying that Dumpy’s presence is required for posterior lobe cells to elongate and project from the lateral plate.

**Figure 7.**
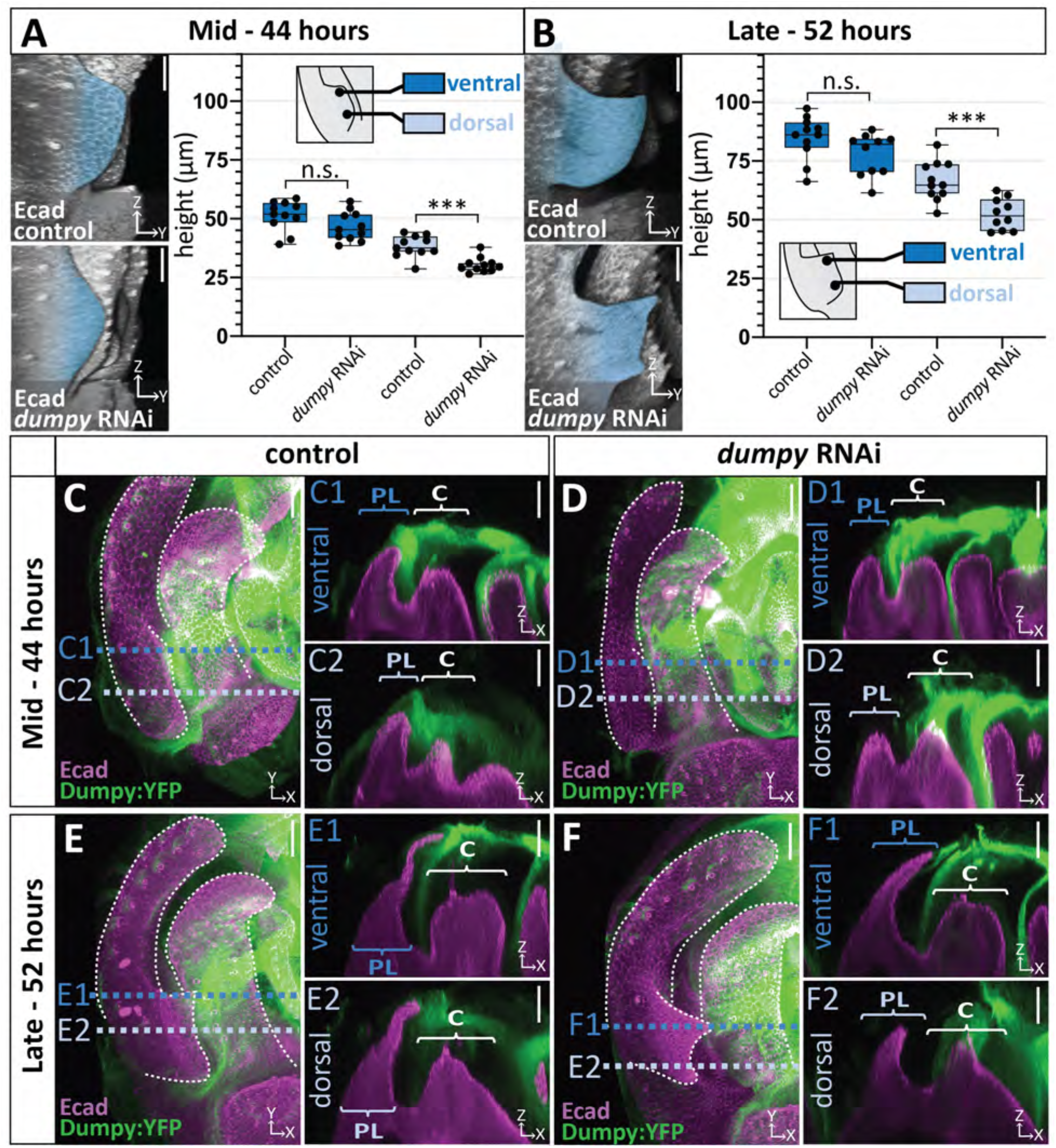
Correlation between the deposition of Dumpy and knockdown phenotype. (A-B) Comparison of *mCherry* RNAi (control) and *dumpy* RNAi at 44 hours APF (A) and 52 hours APF (B). Images are rotated in 3D to visualize the full shape of the posterior lobe labeled with E-cadherin. Quantification of tissue height at the ventral tip (dark blue) and dorsal base (light blue) of the posterior lobe. Cartoon represents relative location of cross-section used for tissue thickness measurement. Individual data points presented; n=at least 10 per each time point. The ventral tip is defined as the location where the posterior lobe is at its max height. The base was determined by moving 19.76μm dorsally from the ventral tip. Statistical significance for each time point indicated (unpaired t-test; ***p≤0.001; n.s.=not significant p≥0.05). (C-F) Comparison of *mCherry* RNAi (control) (C & E) and *dumpy* RNAi (D & F) at 44 hours APF and 52 hours APF with Dumpy:YFP (Green) and Ecad (Magenta). GFP antibody was used to increase YFP signal. All cross-sections are oriented with apical side at the top and basal side at the bottom. Relevant structures labeled: Lateral plate (LP) posterior lobe (PL), and clasper (C). Cross-sections are max projections of 5.434μm sections to show full Dumpy connection. Images were independently brightened to show relevant structures. Scale bar, 20μm. n=at least 5 per experiment.

## Discussion

Here, we determined how a morphological novelty forms at the cellular level, and in doing so, revealed distinctive cell and aECM interactions underlying its development and evolution. We identified how an extreme change in the shape of cells in the developing posterior lobe accounts for its novel morphology. While intrinsic cytoskeletal components may contribute to this process, our results highlight the critical role played by a vast extrinsic network of ECM on the apical side of the epithelium. It was unexpected that such an elaborate supercellular matrix structure would participate in the evolution of a seemingly simple novelty. Below, we consider the potential roles played by the aECM in posterior lobe development and diversification, and discuss how studies of morphogenesis can illuminate the simple origins of structures that might otherwise seem impossibly complex to evolve.

### Mechanisms for aECM-mediated control of cell height in the posterior lobe

Our work demonstrates an important role for the aECM protein, Dumpy, in the growth of the posterior lobe, as exhibited by the dramatic phenotypes in the *dumpy* RNAi background and the strong association of Dumpy:YFP with only the tallest cells in these experiments. Our data is consistent with three possible mechanisms. First, Dumpy could serve as a structural support while autonomous cell mechanical processes drive apico-basal elongation. Second, the cells of the posterior lobe could be pulled mechanically through their connection to the Dumpy aECM. This process could operate passively, deforming cells of the lobe, but could also drive changes in the cytoskeleton in response to external tensions. Finally, the aECM could play a direct role by altering cell signaling dynamics, as has been exhibited by the basal ECM (Kirkpatrick et al., 2004; Kreuger et al., 2004; Wang et al., 2008). Previous research has shown that the JAK/STAT pathway is important for posterior lobe development (Glassford et al., 2015), and their ability to signal to the correct cells could be altered in the absence of Dumpy. Of course, these models are not mutually exclusive and some combination of these mechanisms may be integrated to shape the posterior lobe. Our observations of increased cytoskeletal components in posterior lobe cells and the reduced height of cells that lack Dumpy in our knockdown experiments are consistent with all three mechanisms, which are difficult to differentiate experimentally. When we examine morphogenesis in non-lobed species, we observed that the lateral plate drops below the clasper (Figure 1 - supplement 1). Assuming this ancestral process still occurs in lobed species, it is quite possible that the aECM ‘holds’ cells of the posterior lobe during the early stages of posterior lobe development while the lateral plate is pulled down, causing cells of the posterior lobe to elongate to relieve the stress. Future manipulative biomechanical studies will be required to explore these possibilities.

### The role of aECM in the diversification of genital structures

Genitalia represent some of the most rapidly diversifying structures in the animal kingdom, and our results suggest the aECM may participate in the modification of *Drosophila* genital structures. The shape of the posterior lobe is extremely diverse among species of the *melanogaster* clade (Coyne, 1993). Our results demonstrate that reducing the levels of Dumpy can affect the shape of the posterior lobe, with extreme knockdown phenotypes approximating the posterior lobe of *D. mauritiana*. Furthermore, the clasper and phallus show dense deposits of Dumpy, suggesting that the aECM could play important roles in diversifying these remarkably variable structures. During the course of evolution, one could imagine that by altering which cells are connected to the aECM, the strength of those connections, or the forces acting on those connections could lead to changes in morphological shape. Hence identifying causative genes that differentiate these structures could uncover novel mechanisms for genetically controlling the behavior of this aECM and behaviors of cells bound to this dynamic scaffold.

### Integrating cells into a pre-existing aECM network to generate morphological novelty

In comparing the morphogenesis of a novel structure to close relatives which lack it (representing a proxy for the ancestral state), we identified a likely path by which the aECM became associated with the posterior lobe. The aECM, while understudied, has been implicated in the morphogenesis of many structures (Diaz-de-la-Loza et al., 2018; Etournay et al., 2015; Ray et al., 2015; Fernandes et al., 2010; Dong et al., 2014; Heiman et al., 2009; Low et al., 2019), and yet, its role during the evolution of novel structures is largely unexplored. We find a conserved aECM network associated with central genital structures (clasper, sheath, and phallus) in both lobed and non-lobed species. In non-lobed species, *dumpy* is expressed weakly at the base between the lateral plate and clasper resulting in a thin connection of aECM from clasper to the crevice (Figure 8). By contrast, lobed species express high levels of *dumpy* between the presumptive posterior lobe and clasper, resulting in large amounts of aECM in the crevice. We hypothesize that this increase in aECM allows cells at the base of the lateral plate to be integrated into this ancestral aECM network (Figure 8), a step which was likely significant to the evolution of the posterior lobe. Overall, this suggests that the aECM could be an unexpected target for generating novel anatomical structures.

**Figure 8.**
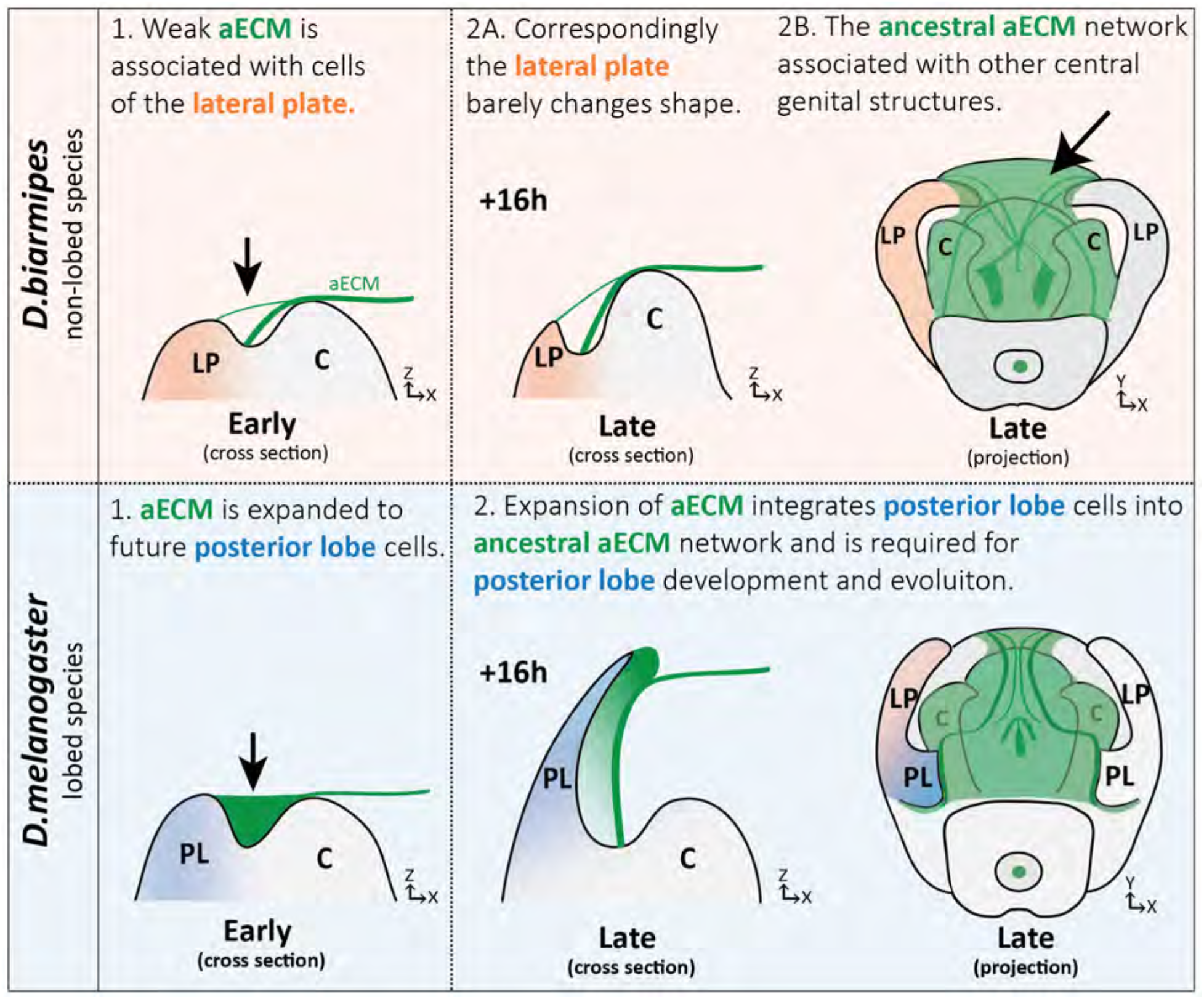
Expansion of apical extracellular matrix underlies the morphogenesis of a recently evolved structure. (Top) Illustration of non-lobed species, *D. biarmipes*, with ancestral aECM network covering central genital structures (2B) including the clasper (C), sheath, and phallus. Weak connections of aECM span from the clasper to the lateral plate (LP) during early development (1 & 2A - top). (Bottom) Illustration of lobed species, *D. melanogaster*. The aECM network has expanded to fill the crevice between the lateral plate and clasper (1-bottom) integrating these cells into the ancestral aECM network (2-bottom). This aECM population is needed for cells to properly project from the lateral plate, forming the posterior lobe.

The expanded *dumpy* expression we observed caused us to consider how the posterior lobe gained this aECM attachment. Interestingly, our previous work found a gene regulatory network (GRN) that regulates development of an ancestral embryonic structure, the posterior spiracles, which was co-opted during the evolution of the posterior lobe and regulates its development (Glassford et al., 2015). Previous work has shown that *dumpy* is expressed in the posterior spiracles (Wilkin et al., 2000), and we have observed a thin tether of Dumpy:YFP connecting the posterior spiracles to the surrounding embryonic cuticle (Figure 8 - Supplement 1). This is consistent with previously identified roles for Dumpy in epithelia-cuticle attachment in the wing (Etournay et al., 2015; Ray et al., 2015) and hypothesized role in the muscle, leg, and antenna <u>(Wilkin et al., 2000; Ray et al., 2015)</u>. Identification of regulatory elements which activate *dumpy* in the posterior lobe will be necessary to determine whether its role in the posterior spiracle was relevant to the evolution of expanded genital expression.

Evolution is thought to act through the path of least resistance. When confronted with the remarkable diversity of genital morphologies present in insects, one must wonder how the intricate projections, bumps, and divots form in its underlying epithelia. Models of co-option have been appealing because they establish pre-existing mechanisms in place that can be rapidly ported to new locations to generate massive changes in a tissue. Our examination of the cellular processes during posterior lobe morphogenesis highlights a different way that co-option may work. Here, the aECM mechanism we uncovered appears to be a path of least resistance because this tissue already uses a vast network of aECM to potentially pattern other structures, such as the phallus and its multiple elaborations (Rice et al., 2019; Peluffo, et al. 2015; Kamimura, 2010). Because this network of aECM represents a pre-existing condition, it is easy to appreciate how cells of the posterior lobe could evolve novel extracellular connections to this network to generate a new protrusion. On the other hand, tissues which lack such an ancestral network may well be less likely to evolve projections through this mechanism. While the aECM is required for this morphogenetic process, we envision that additional networks and processes must be contributing to the full morphogenesis of the posterior lobe. Determining genetic changes which underlie such remarkable cellular responses represents a major looming challenge in evo-devo research (Smith et al., 2018).

## Materials and Methods

### Key resources table

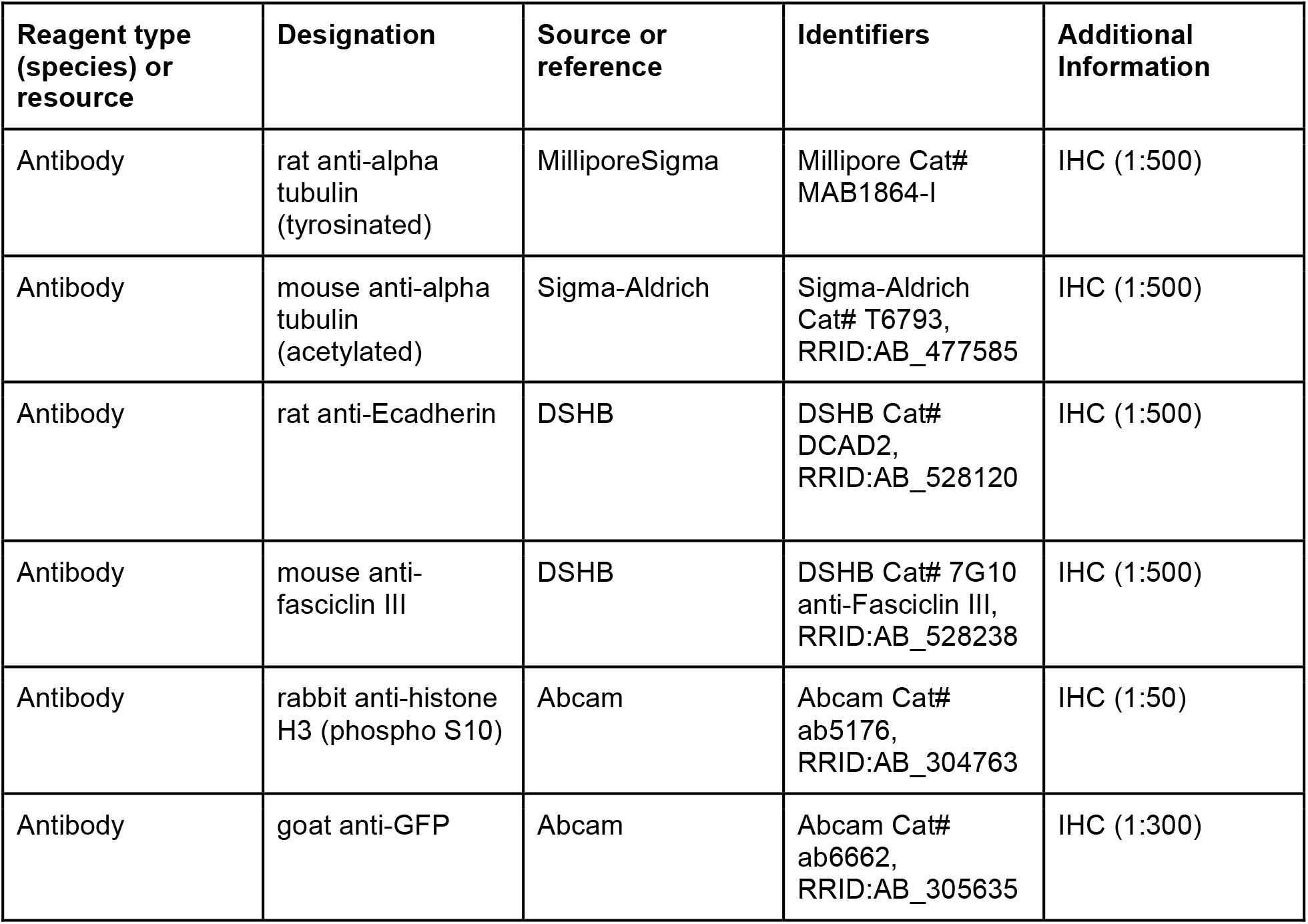
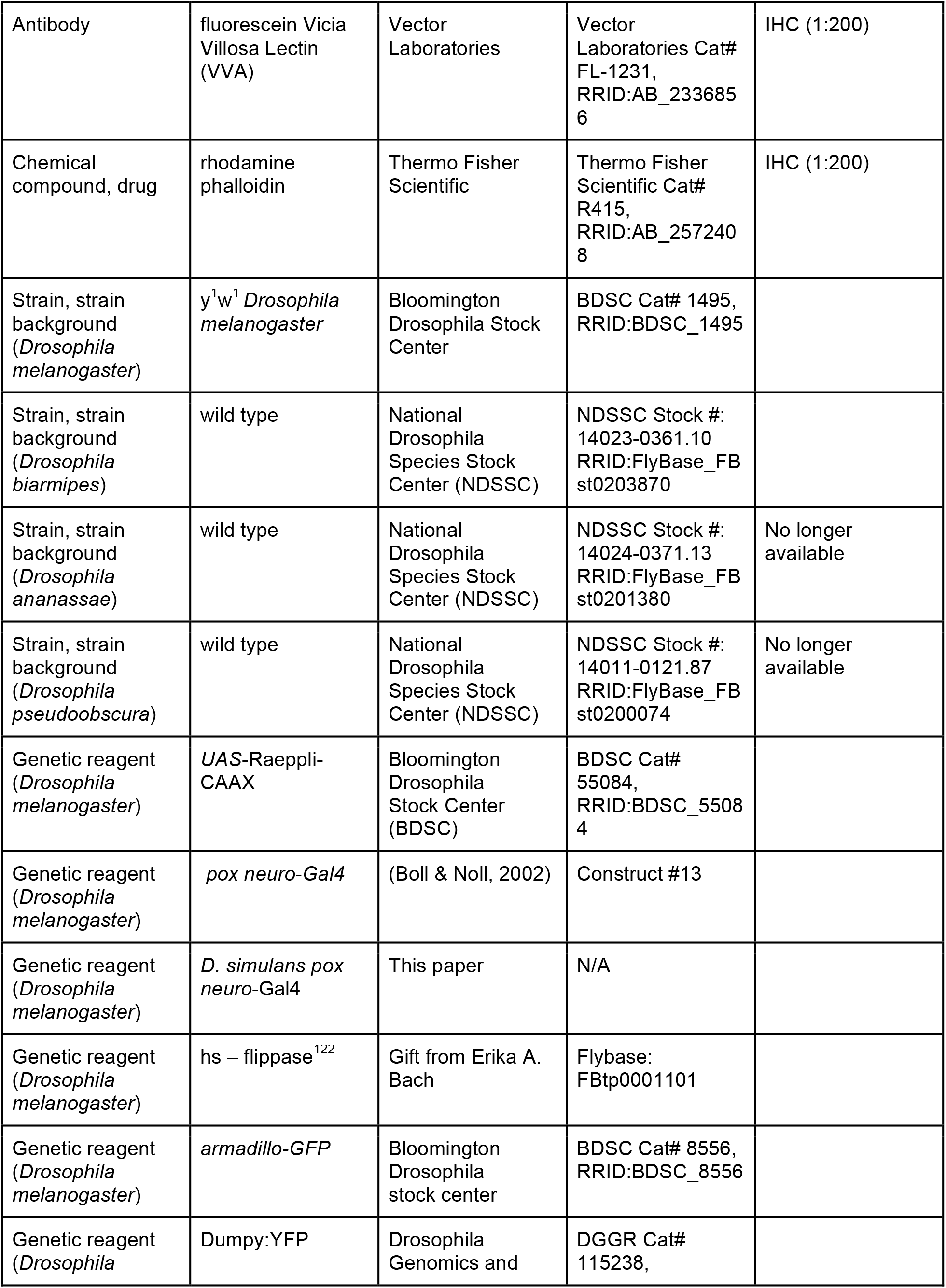
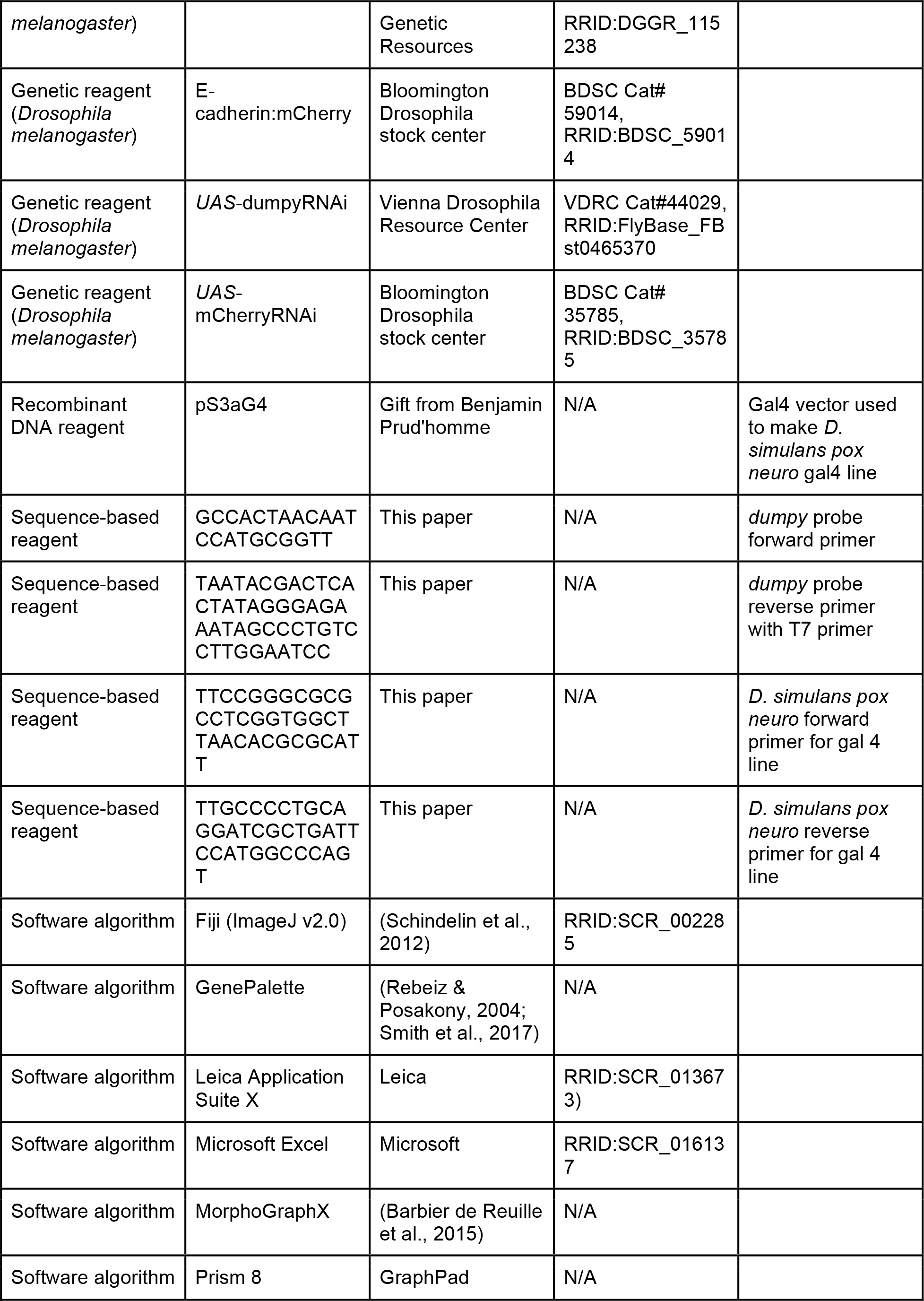

### Fly stocks and genetics

Fly stocks were reared using standard culture conditions. Wild type species used in this study were obtained from the University of California, San Diego *Drosophila* Stock Center (now known as The National Drosophila Species Stock Center at Cornell University)(*Drosophila biarmipes* #14024-0361.10, *Drosophila ananassae* #14024-0371.13, *Drosophila pseudoobscura* #14011-0121.87) and from the Bloomington Drosophila Stock Center (*Drosophila melanogaster* [*y*^1^*w*^1^] #1495). *pox neuro*-Gal4 (construct #13) was obtained from Werner Boll (Boll & Noll, 2002). The following were obtained from the Bloomington Drosophila stock center: *UAS*-Raeppli-CAAX (#55084), *armadillo-GFP* (#8556), Ecadherin:mCherry (#59014), and *UAS*-mCherryRNAi (control for RNAi experiments, as mCherry is not present in the *Drosophila* genome)(35785). *UAS*-dumpyRNAi was obtained from the Vienna Drosophila Resource Center (#44029) and Dumpy:YFP was obtained from the Drosophila Genomics and Genetic Resources (#115238).

For the Raeppli experiments, stable lines of hs-flippase;;*UAS*-Raeppli-CAAX/*UAS*-Raeppli-CAAX and *D. simulans pox neuro*-gal4/*D. simulans pox neuro*-gal4;*UAS*-Raeppli-CAAX/*UAS*-Raeppli-CAAX were generated. *D. simulans pox neuro*-gal4 was used as opposed to *pox neuro*-gal4 because a gal4 driver on the second chromosome was required. Virgin females from the first line were crossed to males from the second line to ensure hs-flippase was inherited by all offspring. Offspring were collected and grown as normal, heat shocked at 37°C for 1 hour around 24 to 28 hours APF, and allowed to finish development at 25°C.

### Sample Preparation

Pupal samples were prepared following the protocol in Glassford, et al., 2015. Briefly, samples were incubated at 25°C unless otherwise noted. Dissections were performed in cold PBS, pupae were cut in half, removed from their pupal cases, and fat bodies removed by flushing. Larval samples were dissected in cold PBS by cutting the larva in half, and flipping the posterior end of the larva inside out. All samples were fixed for 30 minutes at room temperature in PBS with 0.1% Triton-X and 4% paraformaldehyde. Samples stained with phalloidin had Triton-X concentrations increased to 0.3%. Samples used for VVA staining were removed from pupal cuticle before being fixed in PBS with 0.1% Triton-x, 4% paraformaldehyde, and 1% trichloroacetic acid on ice for 1 hour followed by 30 minutes at room temperature. The trichloroacetic acid method causes some slight tissue distortion, as the precipitation treatment utilized to refine the VVA signal causes the posterior lobe to become slightly deformed and curve in towards the clasper. However, similar defects were not observed in the other structures such as the lateral plate or in *D. biarmipes*. Samples were stored in PBT for immunostaining at 4°C for up to two days. For *in situ* hybridization, samples were rinsed twice in methanol and rinsed twice in ethanol. Samples were stored at −20°C in ethanol.

### Immunostaining and *in situ* hybridization

For immunostaining, genital samples were removed from the surrounding pupal cuticle and incubated overnight at 4°C with primary antibodies diluted in PBS with 0.1% Triton-X (PBT). VVA and phalloidin samples were placed on a rocker. The following primary antibodies were used: rat anti-alpha tubulin (tyrosinated) 1:500 (MAB 1864-I, MilliporeSigma), mouse anti-alpha tubulin (acetylated) 1:500 (T6793, Sigma-Aldrich), rat anti-Ecadherin 1:500 (DCAD2, DSHB), mouse anti-fasciclin III 1:500 (7G10, DSHB), rabbit anti-histone H3 (phospho S10) 1:50 (ab5176, Abcam), goat anti-GFP 1:300 (ab6662, Abcam), fluorescein Vicia Villosa Lectin (VVA) 1:200 (FL-1231, Vector Laboratories). The goat anti-GFP was used to increase signal of Dumpy:YFP in the knockdown experiments only. Primary antibody was removed by performing two quick rinses and two long washes (at least 5 minutes) in PBT. Samples were incubated overnight at 4°C in secondary antibodies diluted in PBT. The following secondary antibodies were used: donkey anti-rat Alexa 594 1:500 (A21209, Invitrogen), donkey anti-mouse Alexa 488 1:500 (A21202, Thermo Fisher Scientific), donkey anti-rat Alexa 488 1:500 (A21208, Thermo Fisher Scientific), goat anti-mouse Alexa 594 1:500 (A-11005, Thermo Fisher Scientific), goat anti-rabbit Alexa 594 1:500 (A-11012, Thermo Fisher Scientific), donkey anti-goat Cy2 1:500 (705-225-147, Jackson ImmunoResearch). Rhodamine phalloidin (R415, Thermo Fisher Scientific) stain was performed with secondary antibody. Samples were washed out of secondary antibody by performing two quick rinses and two long washes (at least 5 minutes) in PBT. Samples were then incubated in 50% PBT/50% glycerol solution for at least 5 minutes. Pupal samples were mounted on glass slides coated with Poly-L-Lysine Solution. Glass slides had 1 to 2 layers of double side tape with a well cut out in which the sample was placed and covered with a cover slip

*in situ* hybridization was performed following the protocol in Rebeiz et al., 2009 with modifications to perform *in situs* in the InsituPro VSi robot (Intavis Bioanalytical Instruments) as done by Glassford et al., 2015.

### Microscopy and live imaging

Cuticles of adult posterior lobes and in situ hybridization samples were imaged on Leica DM2000 with a 40x objective for cuticles and a 10x objective for in situ samples. Samples with fluorescent antibodies and fluorescently tagged proteins were imaged using a Leica TCS SP5 Confocal microscope using either a 40x or 63x oil immersion objective.

To live image genital development, a 2% agar solution was poured into a small petri dish filling the dish half way. A 0.1-10μL pipette tip was used to make small wells in the agar for pupal samples. Timed pupal samples were inserted head first into the small well and a 5-300μL pipette tip was used to push sample into agar by placing the tip around the posterior spiracles on the pupal case. To better image the developing genitalia the pupal case at the posterior end was removed with forceps. Deionized water was used to cover the samples and imaged on a Leica TCS SP5 Confocal microscope using a 63x water objective.

To live image embryos, Dumpy:YFP flies were grown in egg-laying chamber with grape agar plates (Genesee Scientific). Embryos were removed from plates using forceps and rolled on a piece of double sided tape to remove the chorion. Embryos then were positioned on a glass coverslip coated with embryo glue. A glass slide was covered with double sided tape and a well was made and filled with halocarbon 27 oil. The cover slip with the embryos was then placed on the glass slide, submerging the embryos in halocarbon oil. Embryos were imaged on a Leica TCS SP8 confocal with a 63x oil objective.

### Image analysis

Images were processed with Fiji (Schindelin et al., 2012) and Photoshop. Three-dimensional views were completed in MorphoGraphX (Barbier de Reuille et al., 2015) or Leica Application Suite X. Movies were processed in Fiji and cell rearrangements were tracked using the manual tracking plugin. Tissue thickness/cell height during development was measured in cross-section view by drawing a line centered between the two sides (based on apical membrane) of the lobe until the basal side was reached. Area of adult posterior lobe cuticles and height of the adult lobe were measured by using the lateral plate as a guide for determining the bottom boundary of the posterior lobe. To prevent any possible bias for one lobe vs the other (i.e. left vs right) which lobe was used in statistical analysis was randomly decided, except for Figure 6 - supplement 1 where both sides of the posterior lobe were considered.

### Transgenic Constructs

To make the *D. simulans pox neuro*-gal4 driver, the posterior lobe enhancer for *pox neuro* in *D. simulans*, identified in Glassford et al., 2015, was cloned using primers listed in key resources table using genomic DNA purified with the DNeasy Blood and Tissue Kit (QIAGEN). Primers were designed using sequence conservation with the GenePalette software tool <u>(Rebeiz and Posakony 2004; Smith et al., 2017)</u>. The cloned sequence was inserted into the pS3aG4 (Gal4) using *AscI* and *SbfI* restriction sites. The final construct was inserted into the 51D landing site on the second chromosome <u>(Bischof et al., 2007)</u>.

## Acknowledgements

The authors thank the members of the M.R. laboratory for comments and discussion on the manuscript. We thank Werner Boll and Markus Noll for the *pox neuro*-gal4 line, the Bloomington stock center, VDRC, and DGGR stock centers for fly stocks, Benjamin Prud’homme for the s3aG4 vector, Erika A, Bach for the hs-flippase line, and Winslow Johnson for the *D. simulans pox neuro*-gal4 line. This work was supported by the National Institutes of Health (GM107387 to M.R. and HD044750 to L.A.D.).

## Author details

Sarah Jacquelyn Smith

Department of Biological Sciences, University of Pittsburgh

Contributions: Conceptualization, Methodology, Validation, Formal analysis, Investigation, Data curation, Writing-original draft, Writing-review and editing, Visualization

Competing interests: No competing interests declared

ORCID: 0000-0002-1469-1821

Lance A. Davidson

Department of Bioengineering University of Pittsburgh

Contributions: Conceptualization, Methodology, Writing-review and editing, Funding acquisition

Competing interests: No competing interests declared

ORCID: 0000-0002-2956-0437

Mark Rebeiz

Department of Biological Sciences, University of Pittsburgh

Contributions: Conceptualization, Methodology, Writing-review and editing, Supervision, Funding acquisition

Competing interests: No competing interests declared

ORCID: 0000-0001-5731-5570

---

**Supplement 1 - figure 1.**
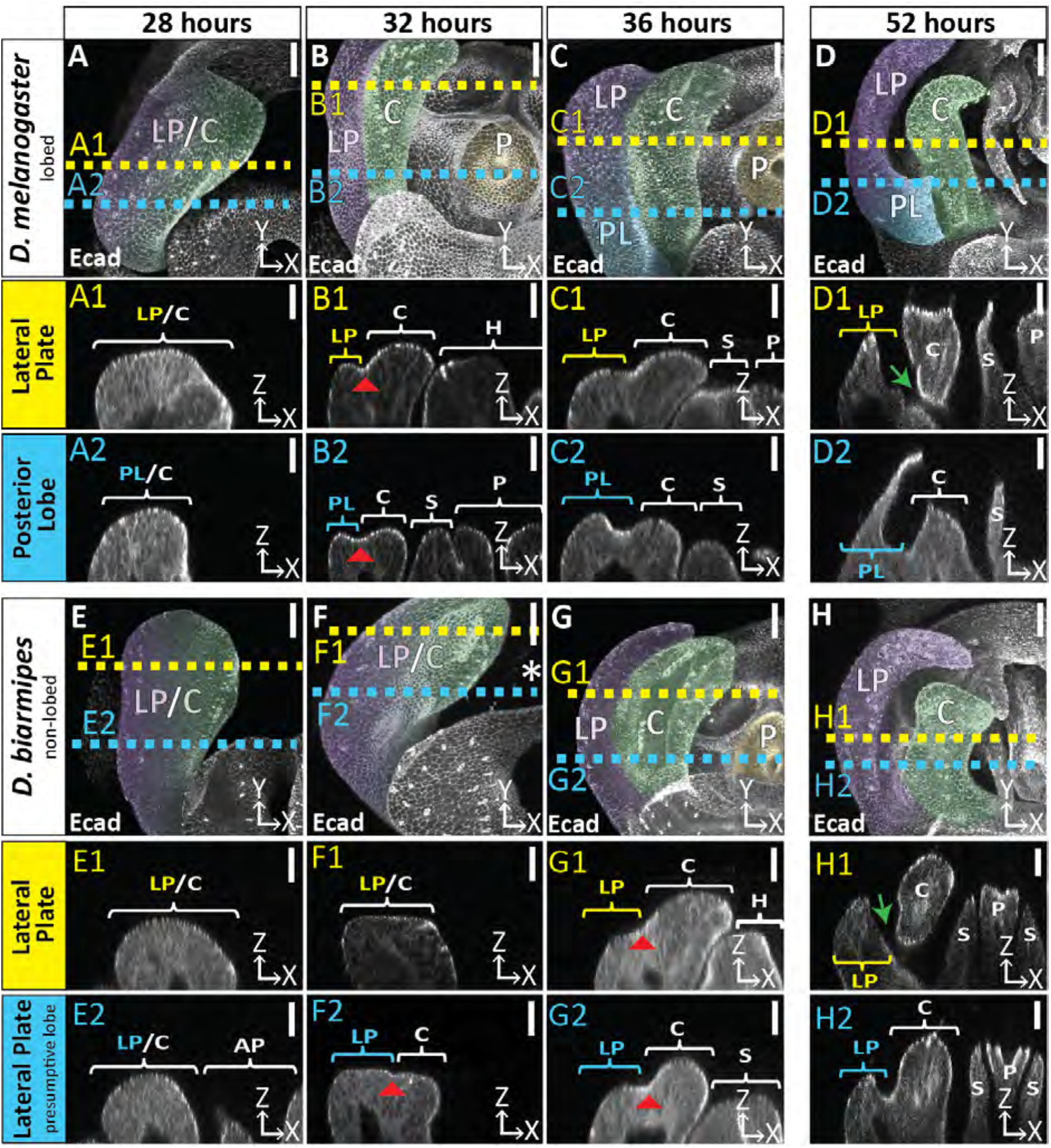
Developmental timing of lobed vs non-lobed genitalia. Developmental time course of the lobed species *D. melanogaster* (A-D) and the non-lobed species *D. biarmipes* (E-H) with E-cadherin label. Location of respective cross sections indicated in yellow for lateral plate and blue for posterior lobe (*D. melanogaster)* or equivalent location in non-lobed species (*D. biarmipes*). Relevant structures are labeled: posterior lobe (PL), lateral plate (LP), clasper (C), sheath (S), and phallus (P). Scale bar, 20μm. At 28 hours APF the genitalia looks relatively similar between *D. melanogaster* (A-A2) and *D. biarmipes* (E-E2). At 32 hours APF in *D. melanogaster* the clasper and lateral plate have fully begun to cleave (B1-2 red arrowhead=cleavage), the lateral plate is lower than the clasper (B1), and the hypandrium, sheath, and phallus have fully everted and are neighboring the clasper and lateral plate (B1-2). *D. biarmipes* lags behind approximately 4 hours. At 32 hours APF there is slight cleavage near the dorsal side of the lateral plate and clasper (F2 red arrowhead), but no cleavage has occurred at the ventral side (F1). In addition, the sheath, hypandrium, and phallus have not everted yet (F1-2). At 36 hours APF in *D. biarmipes*, cleavage has begun along the full length of the lateral plate and clasper (G1-2 red arrowhead), the lateral plate is lower than the clasper (G1-2), and the hypandrium, sheath, and phallus have everted and are next to the lateral plate and clasper (G1-2). As development proceeds later at 52 hours APF the lateral plate and clasper fully separate at the ventral side of the genitalia in both *D. melanogaster* (D1 green arrow) and *D. biarmipes* (H1 green arrow). Full cleavage does not span the length of the lateral plate and clasper (D2 and H2) and stops right before the posterior lobe forms (D2) and also stops before reaching the very dorsal side of the lateral plate and clasper in *D. biarmipes* (H2).

*See supporting file.*

**Supplement 1 - video 1. The posterior lobe protrudes from the lateral plate.** Three-dimensional projections of *D. biarmipes* (left) and *D. melanogaster* (right) samples at 52 hours APF labeled with E-cadherin.

**Figure 2 - supplement 1.**
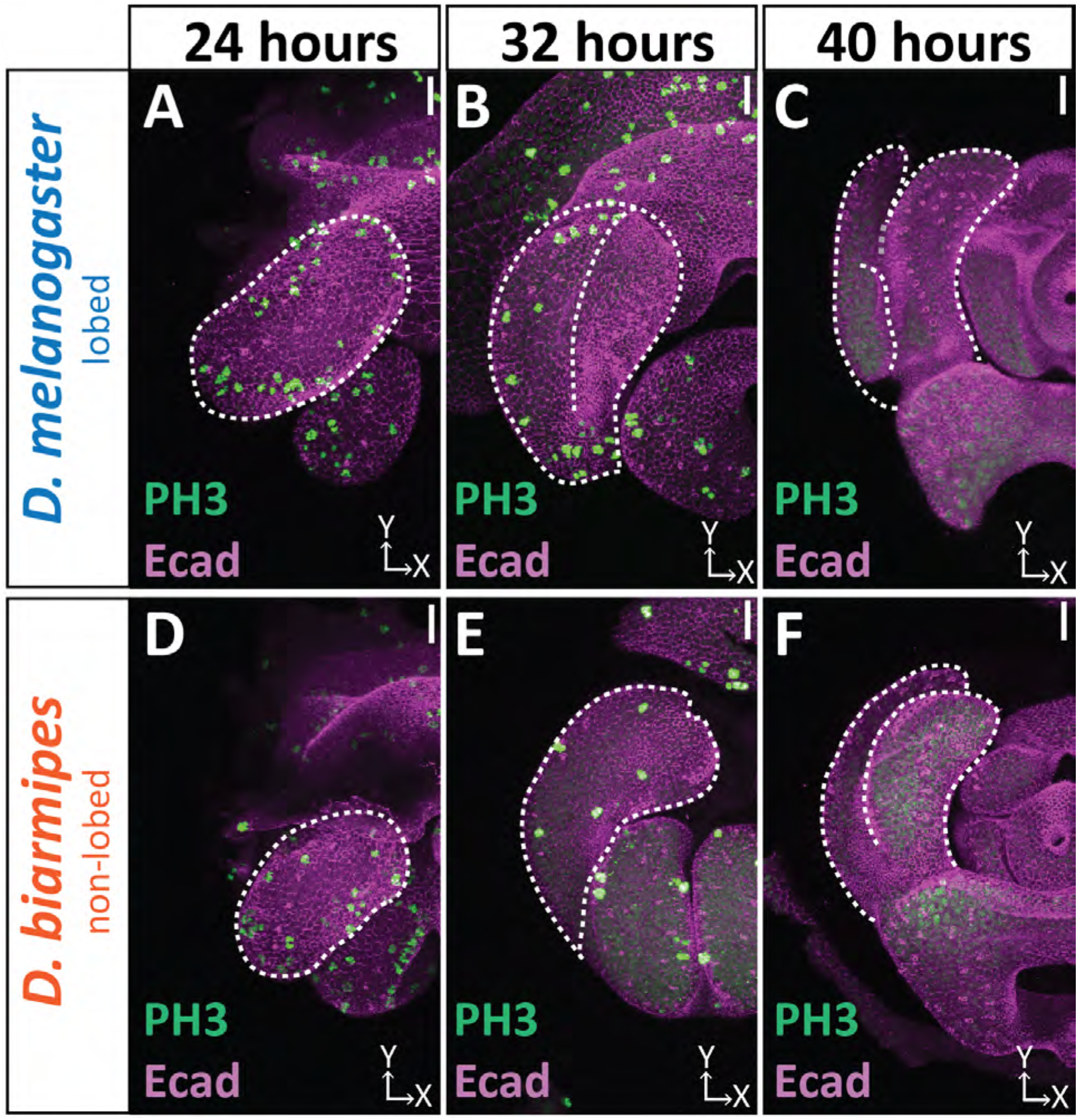
Cell division dynamics do not differ between lobed and non-lobed species. Developmental time course with Phospho-Histone H3 (Ser 10) (PH3; green) labeling actively dividing cells and Ecad (magenta) labeling the apical membrane of the tissue. Only superficial slices are shown to avoid fat body signals beneath lateral plate and clasper. n ≥ 3 per each time point. Scale bar, 20μm. In both *D. melanogaster* and *D. biarmipes* cell division is widespread at 24 hours APF (A & D). Cell division is decreased by 32 hours APF (B & E). By 40 hours APF no cell division is occurring (C & F).

**Figure 2 - supplement 2.**
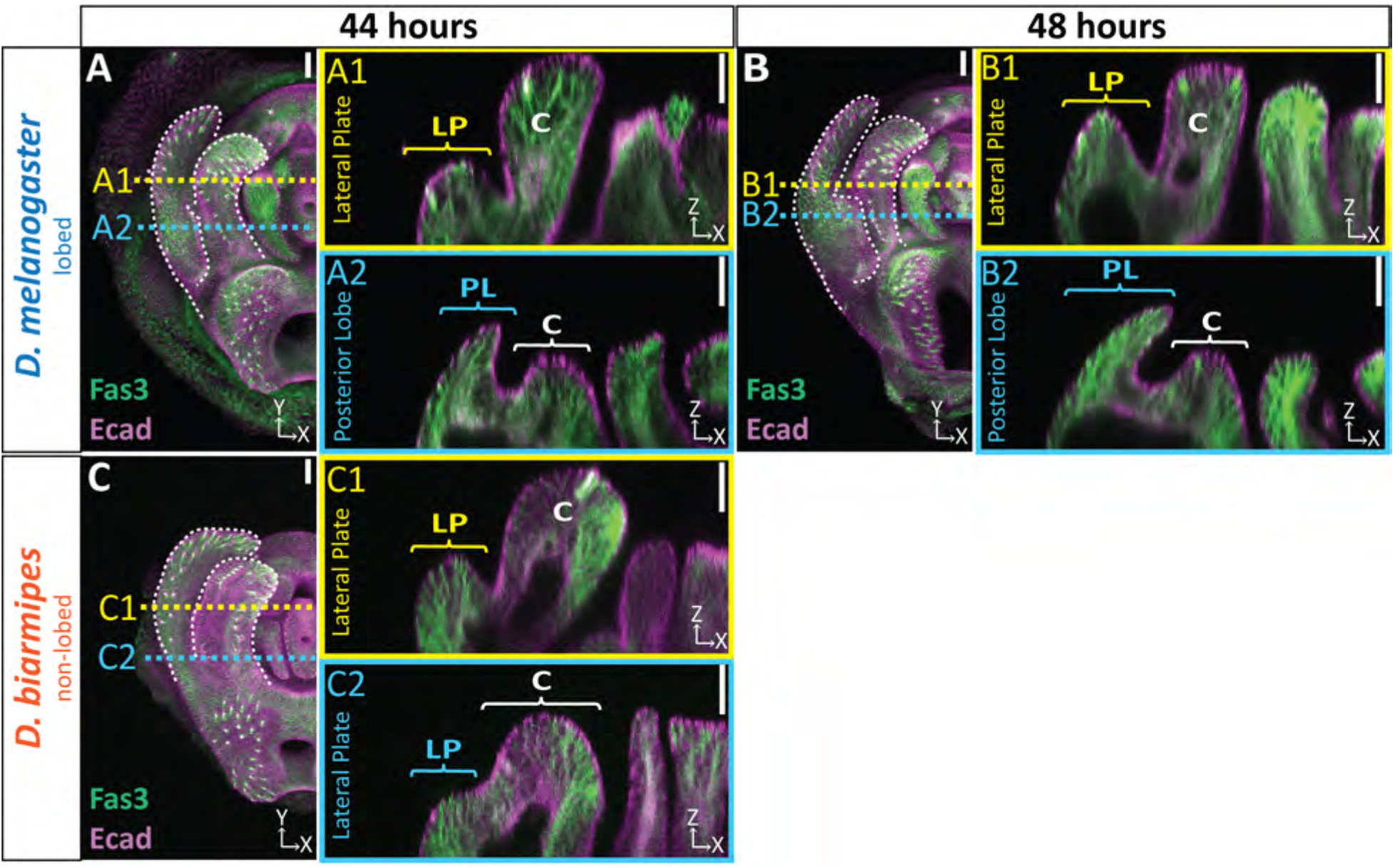
Extended time course of tissue thickness in lobed and non-lobed species. Extended time course for samples quantified in Figure 2F-G. (A-C) Max and cross-section view of 44 hours APF (A & C) and 48 hours APF (B) genital samples with lateral membrane labeled with Fas3 (green) and apical membrane labeled with Ecad (magenta). Location of respective cross sections indicated in yellow for lateral plate (A1-C1) and blue for posterior lobe (*D. melanogaster)* or equivalent location in non-lobed species (*D. biarmipes*) (A2-C2). n≥ 9 per experiment. Scale bar, 20μm.

*See supporting file.*

**Figure 2 - video 1. Minor Cell rearrangement during posterior lobe development.** Live imaging of posterior lobe development with GFP tagged armadillo (apical membrane marker) illustrating a cell dropping from the apical surface and a neighboring cell filling in the gap. Imaging starts at approximately 36 hours APF. Due to uncontrolled temperatures during imaging that were cooler than normal growing conditions, the posterior lobe develops slower and the time indicated is not comparable to other images in the manuscript which were all grown under controlled settings. Based on the thickness of the posterior lobe at the end of the movie the posterior lobe is between 48 to 52 hours APF. Cells were tracked manually and indicated with colored dots. Some dots disappear towards the end of the movie as they become difficult to track due to the signal from cells on the medial side of the posterior lobe.

**Figure 3 - supplement 1.**
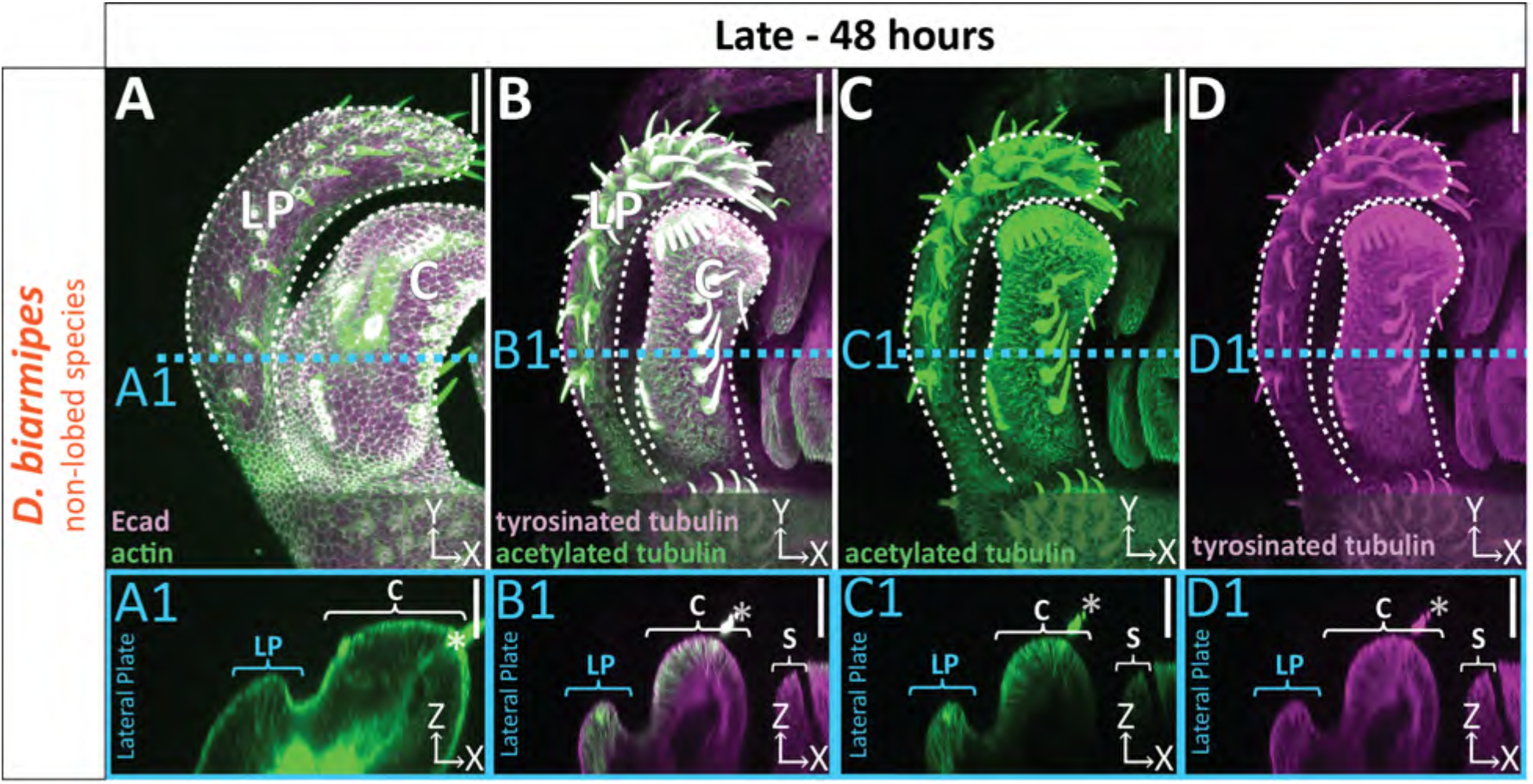
Uniform distribution of cytoskeletal components in non-lobed species. (A-D) Max projections of late (48h APF) genital samples of non-lobed species *D. biarmipes* labeled with F-actin/phalloidin and Ecad (A), acetylated tubulin (B,C), and tyrosinated tubulin (B,D). Location of respective cross sections indicated in blue for presumptive posterior lobe cells (A1-D1). Cross-sections are maximum projection of a restricted 5.434μm thick section to display the full view of the cytoskeleton along the apico-basal axis. All cross-sections are oriented with apical side at the top and basal side at the bottom. Asterisk identifies bristles which have high levels of F-actin and tubulin. Bright basal signal in A1 are fat bodies. Bottom layers were removed in panel A to avoid fat body signal which masked other details. Panels C and D show separate channels of panel B. Relevant structures labeled: Lateral plate (LP), clasper (C), and sheath (S) labeled. Scale bar, 20μm. n≥ 3 per experiment.

**Figure 4 - supplement 1.**
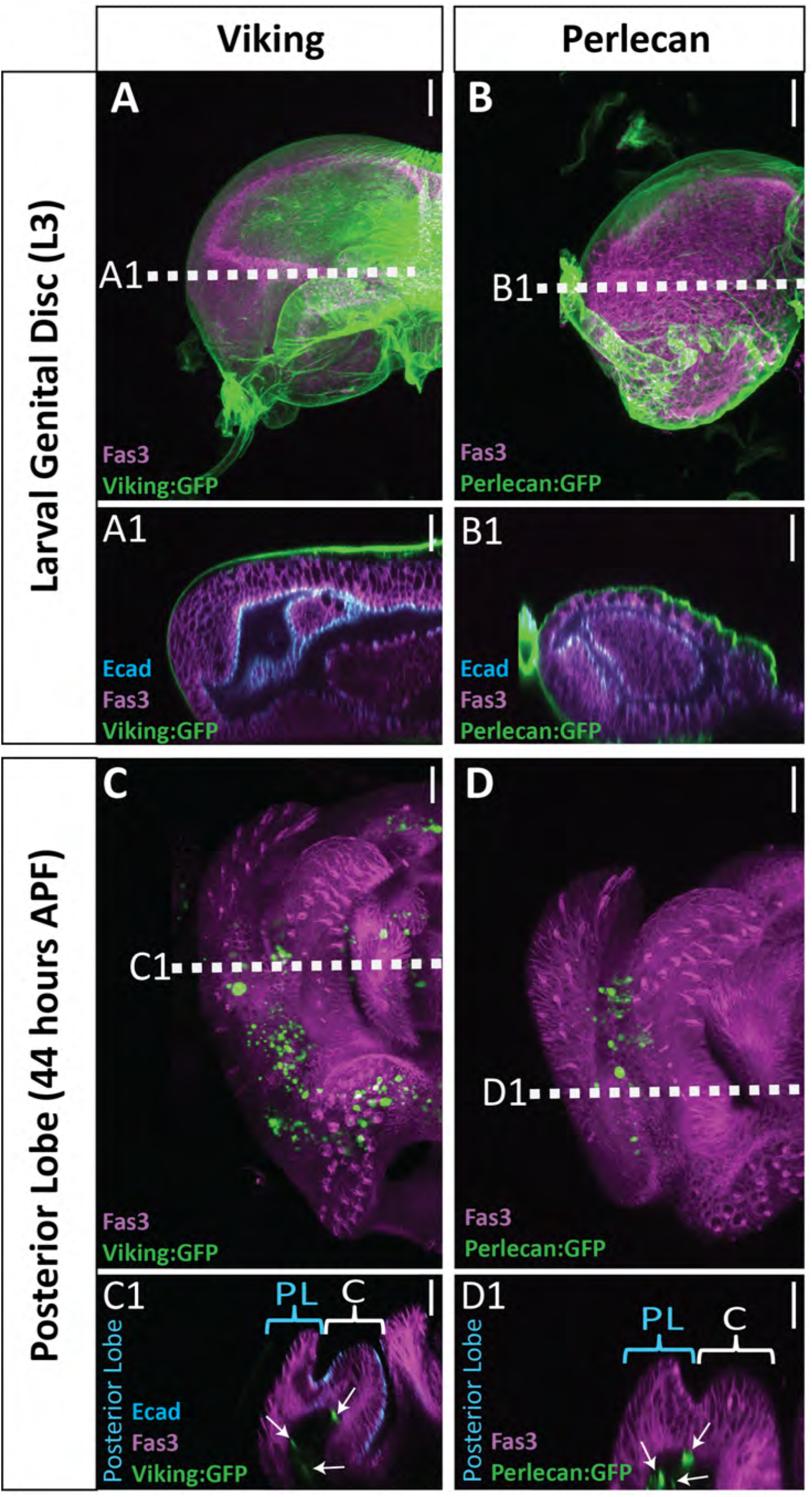
Limited basal ECM present during posterior lobe morphogenesis. Basal ECM markers Collagen IV (Viking:GFP; green)(A & C) and Perlecan (Perlecan:GFP; green) (B & D) in L3 larval genital disc (A & B) and in 44 hours APF genitalia (C & D). Image settings were the same for each marker between larval and pupal samples. Sporadic dots observed are fat bodies (white arrows in cross section), which fill the basal lumen of the pupal genital epithelium. Location of respective cross sections indicated in white. Cross-sections for larval samples are oriented basal sides out, as the disc has not yet everted. Pupal samples are oriented with apical side at the top and basal side at the bottom. Higher amounts of basal ECM are observed in larvae compared to 44 hour APF genital samples. Relevant structures labeled: Posterior lobe (PL) and clasper (C). Scale bar, 20μm.

**Figure 4 - supplement 2.**
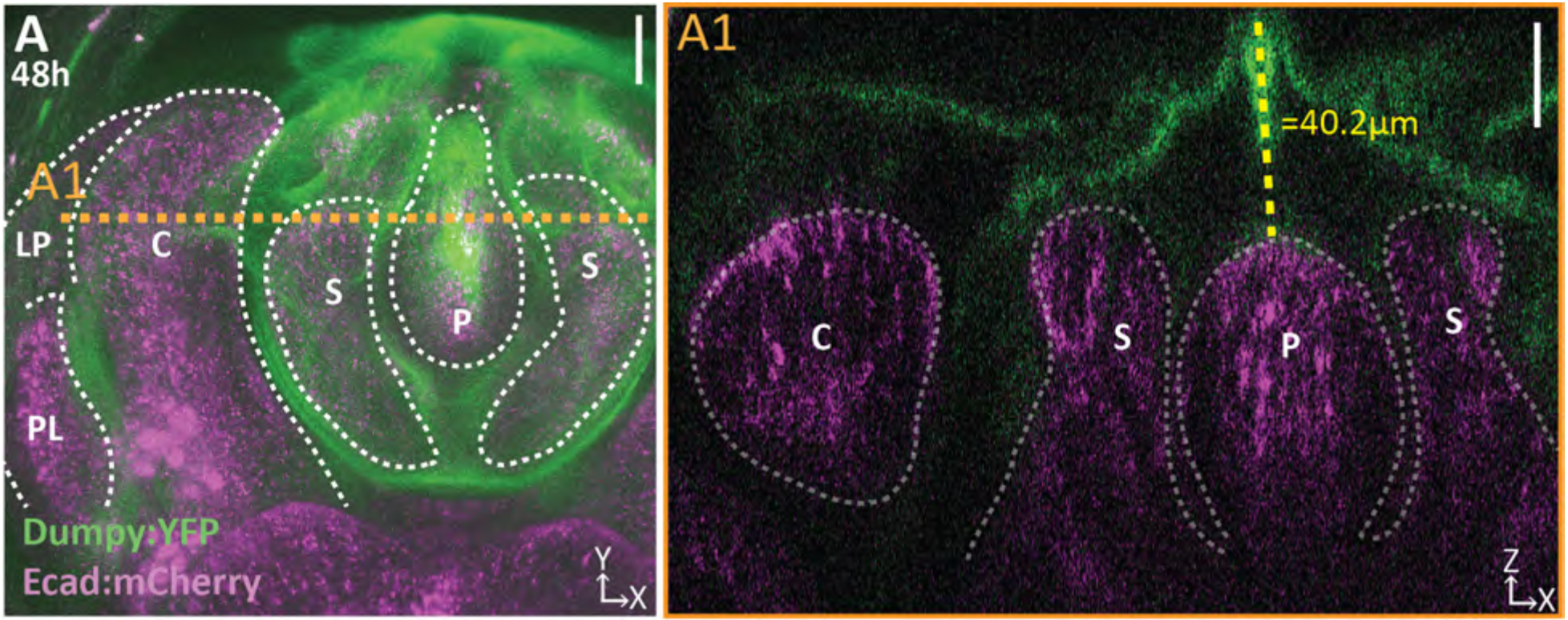
Dumpy extends above the apical surface of the phallus. (A) Projection of Dumpy:YFP (green) and Ecad:mCherry (magenta) imaged live at 48 hours APF. Location of respective cross sections indicated in orange. (A1) Cross section showing extent of Dumpy:YFP observed above the surface of the genitalia. Relevant structures labeled: Posterior lobe (PL), lateral plate (LP), clasper (C), sheath (S), and phallus (P). Scale bar, 20μm. n=3.

**Figure 4 - supplement 3.**
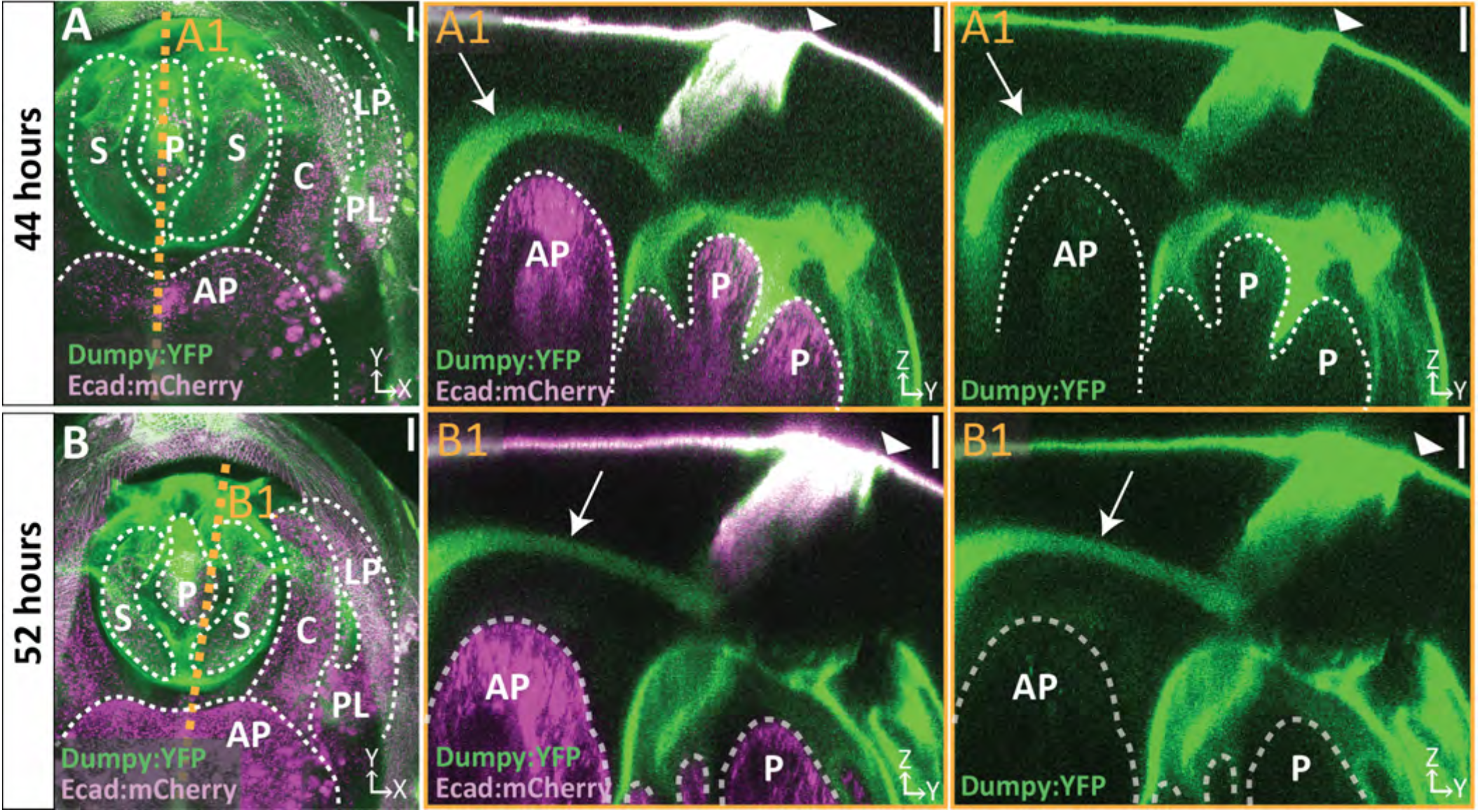
A tether of Dumpy connects the genitalia to the pupal cuticle membrane that encases the developing pupa. (A-B) Live imaging of Dumpy:YFP (green) and Ecad:mCherry (magenta) at respective time points. Location of respective cross sections indicated in orange. (A1-B1) Cross-sections are max projection of a 4.94μm (A1) and 1.73μm (B1) thick section to show full tether (arrow) and its connection to the cuticle (arrowhead) and anal plate. All cross-sections are oriented with apical side at the top and basal side at the bottom. Relevant structures labeled: Posterior lobe (PL), lateral plate (LP), clasper (C), sheath (S), phallus (P), and anal plate (AP). Scale bar, 20μm. n=1 per each time point.

**Figure 4 - supplement 4.**
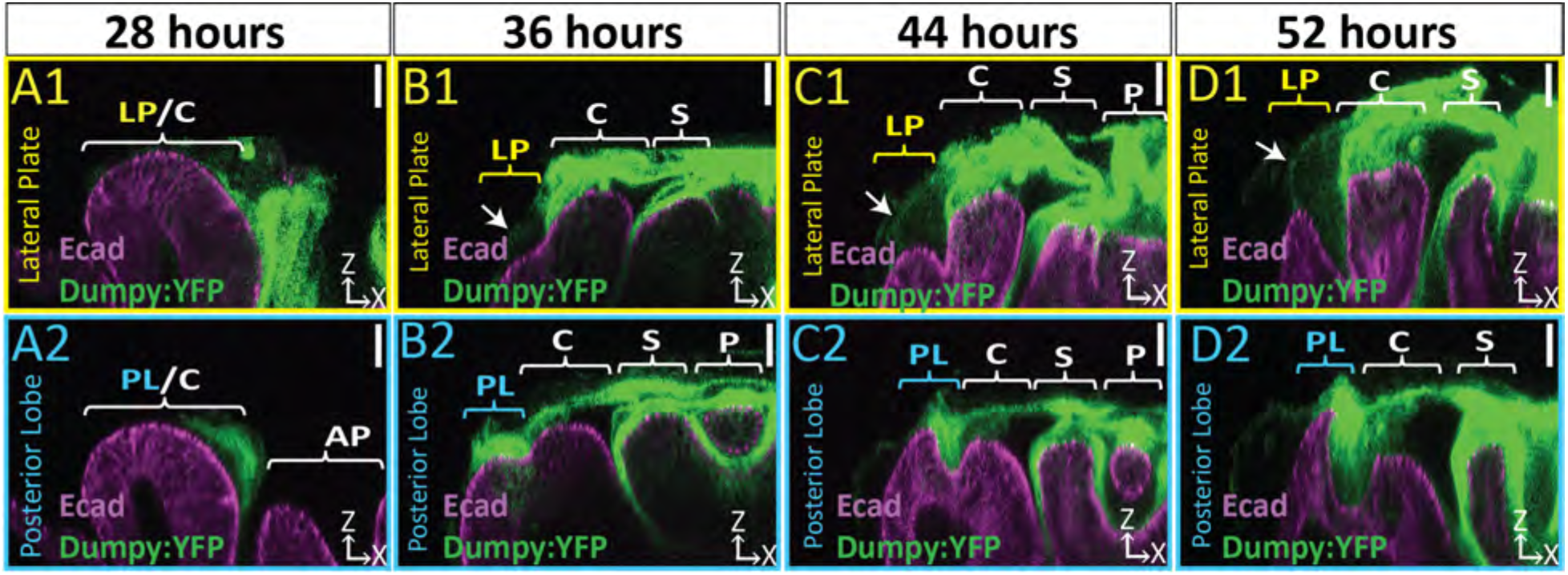
The Dumpy aECM network extends weak connection to lateral plate. (A-D) Brightened images of respective cross sections from Figure 2 of lateral plate (A1-D1) in yellow and posterior lobe in blue (A2-D2). Cross-sections are oriented with apical side at the top and basal side at the bottom. Relevant structure labeled: Posterior lobe (PL), lateral plate (LP), clasper (C), sheath (S), and phallus (P). Scale bar, 20μm. n≥ 4 per experiment. Images were overexposed to show relevant structures.

*See supporting file.*

**Figure 4 - video 1. Three-dimensional structure of the genital Dumpy aECM network.** A 52 hour APF genital sample with Dumpy:YFP (green) and E-cadherin (magenta) is shown. Part 1 of the movie shows a cross-sectional view starting at the ventral side of the posterior lobe and moving towards the dorsal side of the posterior lobe and part 2 shows the same view but starting at the ventral tip of the lateral plate and moving towards the ventral side of the posterior lobe. In the upper-right corner there is a guide that roughly depicts the running location of the cross section. Cross-sections are oriented with apical side at the top and basal side at the bottom. Relevant structures labeled: Posterior lobe (PL), lateral plate (LP), clasper (C), sheath (S), and phallus (P).

*See supporting file.*

**Figure 4 – video 2. A tether of Dumpy connects the genitalia to the surrounding cuticle.** 3D rotation of Dumpy:YFP (green) and Ecad:mCherry (magenta) imaged live at 44 hours APF. Relevant structures labeled: Posterior lobe (PL), lateral plate (LP), clasper (C), sheath (S), phallus (P), and anal plate (AP).

**Figure 5 - supplement 1.**
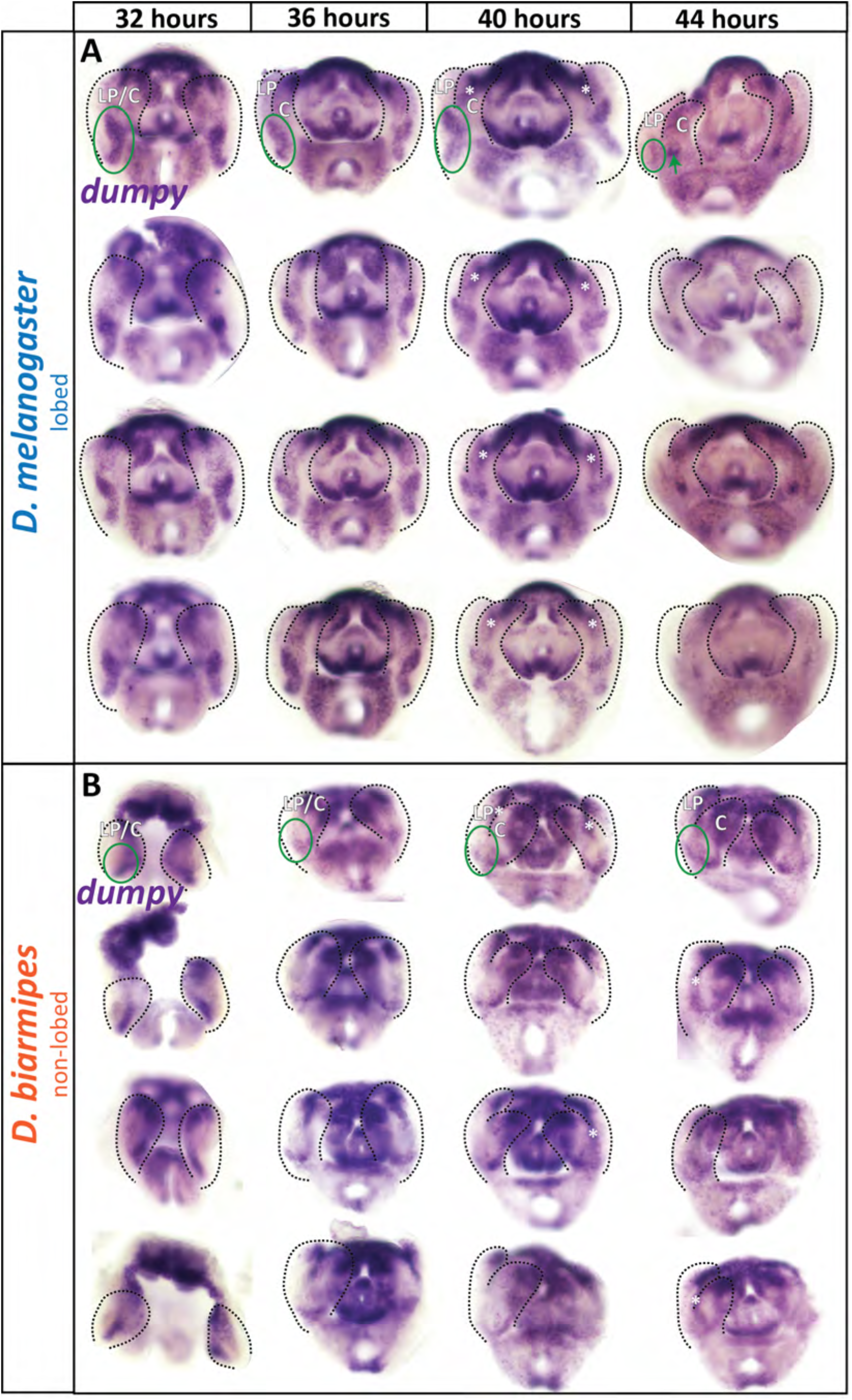
dumpy expression is spatially expanded in lobed species compared to non-lobed species. (A-B) Additional *in situ* hybridization samples for *dumpy* mRNA in lobed species *D. melanogaster* (A) and non-lobed species *D. biarmipes* (B) to show full range of expression observed in experiment. Samples without outlines on one side are due to the tissue being damaged on that side. Green circle in first image highlights relevant location at the base of the lateral plate, but not included in the remaining images to leave images unobstructed. Asterisk indicates the expression is deep in the sample and not expressed in lateral plate or clasper cells. n= 4 per experiment.

**Figure 5 - supplement 2.**
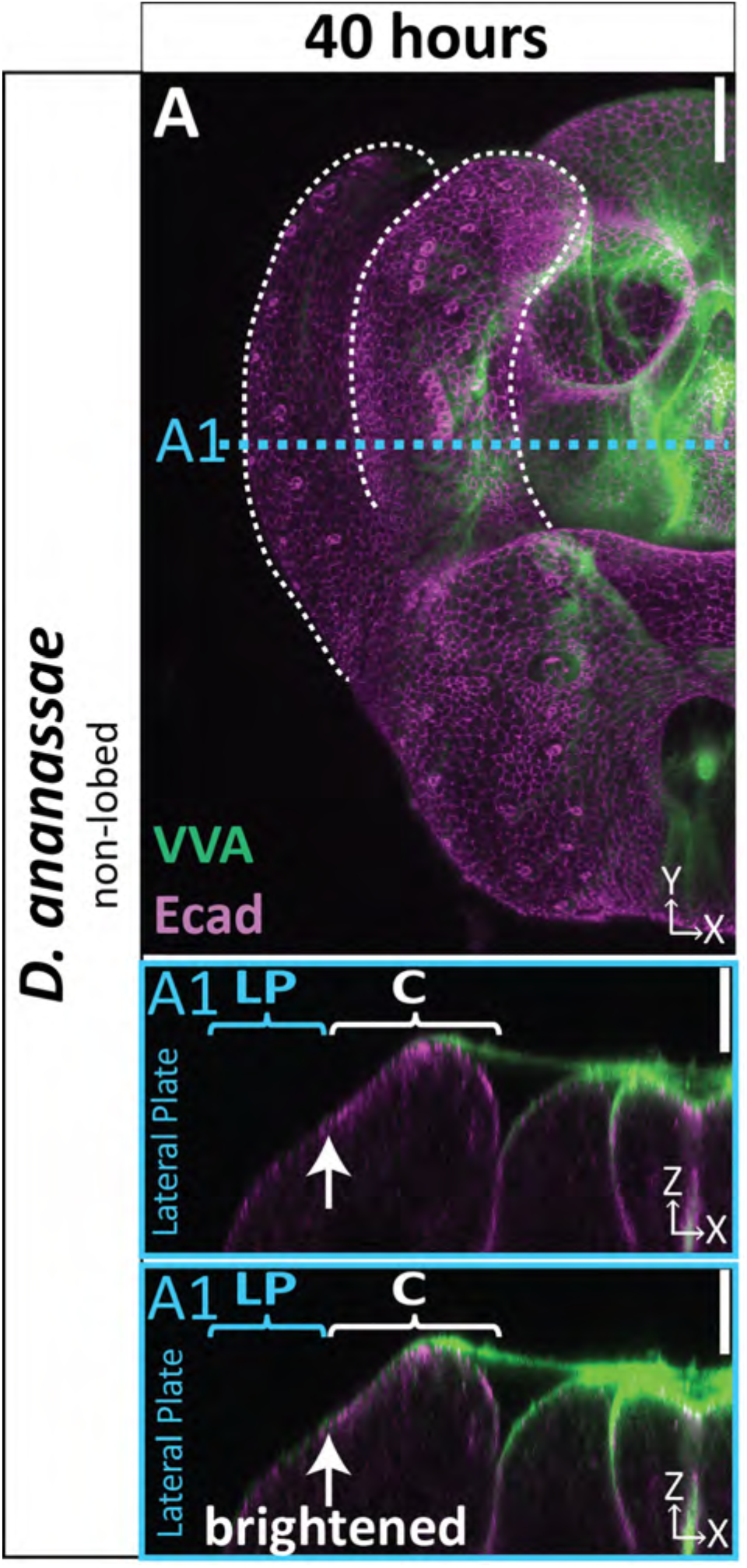
aECM not expanded in non-lobed species *D. ananassae*. (A-C) aECM labeled with VVA (green) and apical membrane labeled with Ecad (magenta) at 40 hours APF in non-lobed species *D. ananassae*. Location of respective cross-sections indicated in blue. Top cross-section displayed with normal brightness to show details and bottom cross-section has been brightened to show where all populations of aECM are located. All cross-sections are oriented with apical side at the top and basal side at the bottom. White arrow highlights the ‘crevice’ between the lateral plate and clasper, which is not pronounced at 40 hours APF in *D. ananassae*. Relevant structures labeled: lateral plate (LP) and clasper (C) labeled. Scale bar, 20μm. n=at least 2 per experiment.

**Figure 6 - supplement 1.**
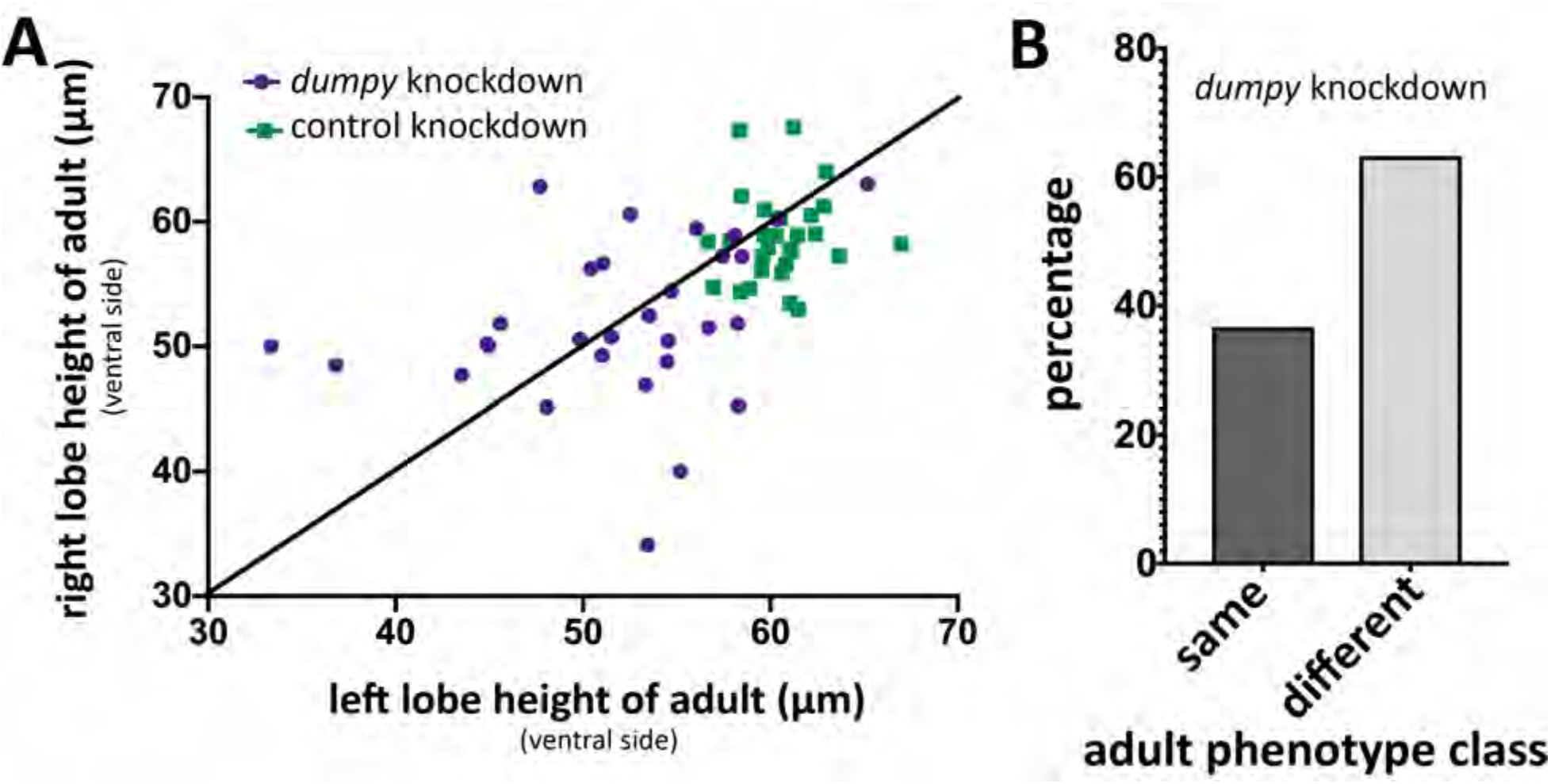
Increased left-right variability of posterior lobe phenotype upon *dumpy* knockdown. (A) Comparison of *dumpy* knockdown (purple circles) and control knockdown (green squares) of left and right adult posterior lobes in single individuals grown at 29°C measuring height at the ventral side of the posterior lobe (single individual represented as a single dot or square). Black line represents perfect correlation in height. *dumpy* knockdown individuals stray more form perfect correlation, indicating that the height of the posterior lobe varies more in the *dumpy* knockdown. (B) Percentage of *dumpy* knockdown individuals plotted in (A) in which both posterior lobes were classified as the same phenotype or different phenotypes (defined in Figure 6).

**Figure 7 - supplement 1.**
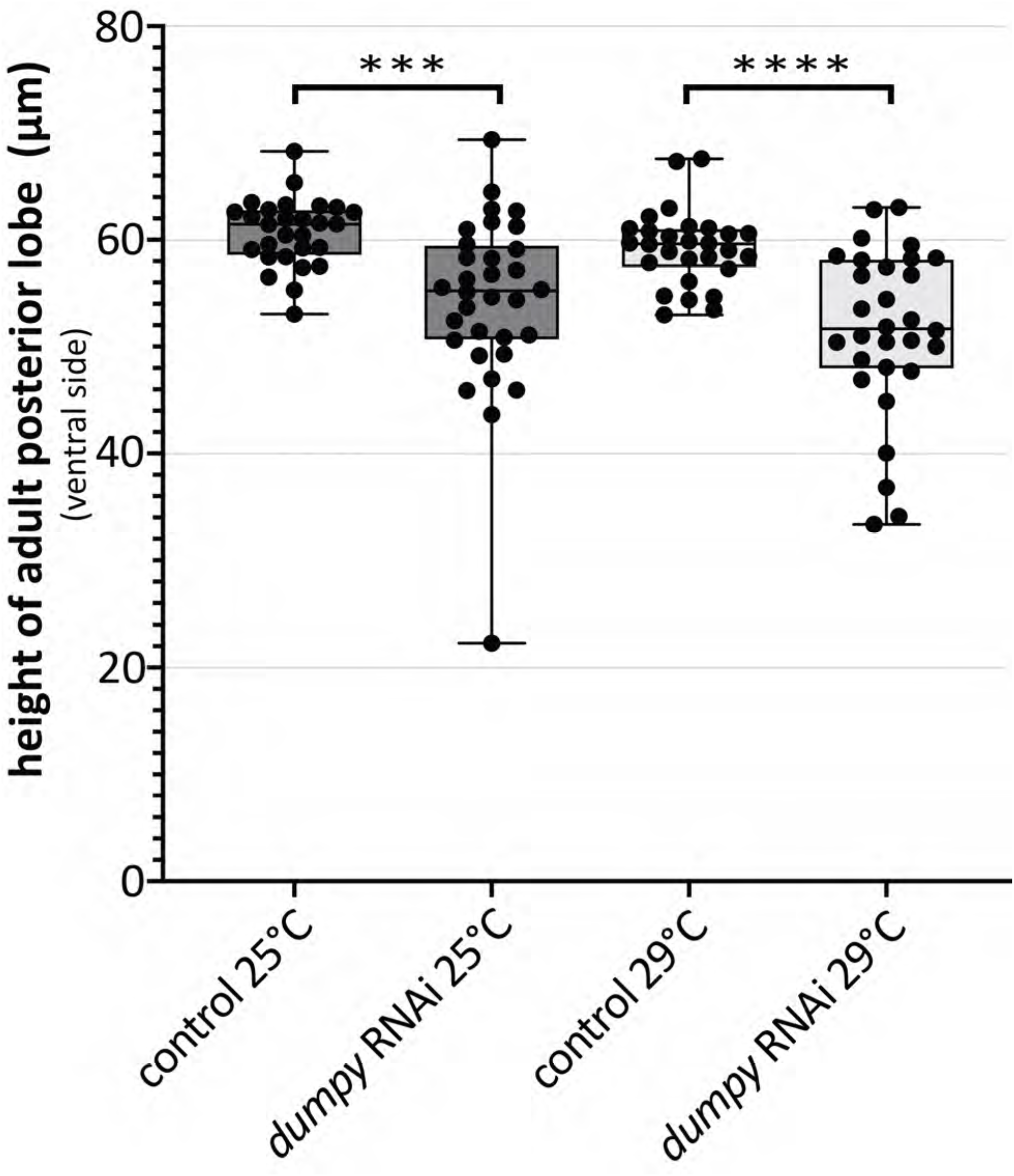
Variability in height of the adult posterior lobe in *dumpy* knockdown. Comparison of *mCherry* RNAi (control) and *dumpy* RNAi adults. Quantification of height of cuticle at the ventral side of the posterior lobe. (unpaired t-test; ***p≤0.001; ****p≤0.0001; n≥28).

**Figure 7 - supplement 2.**
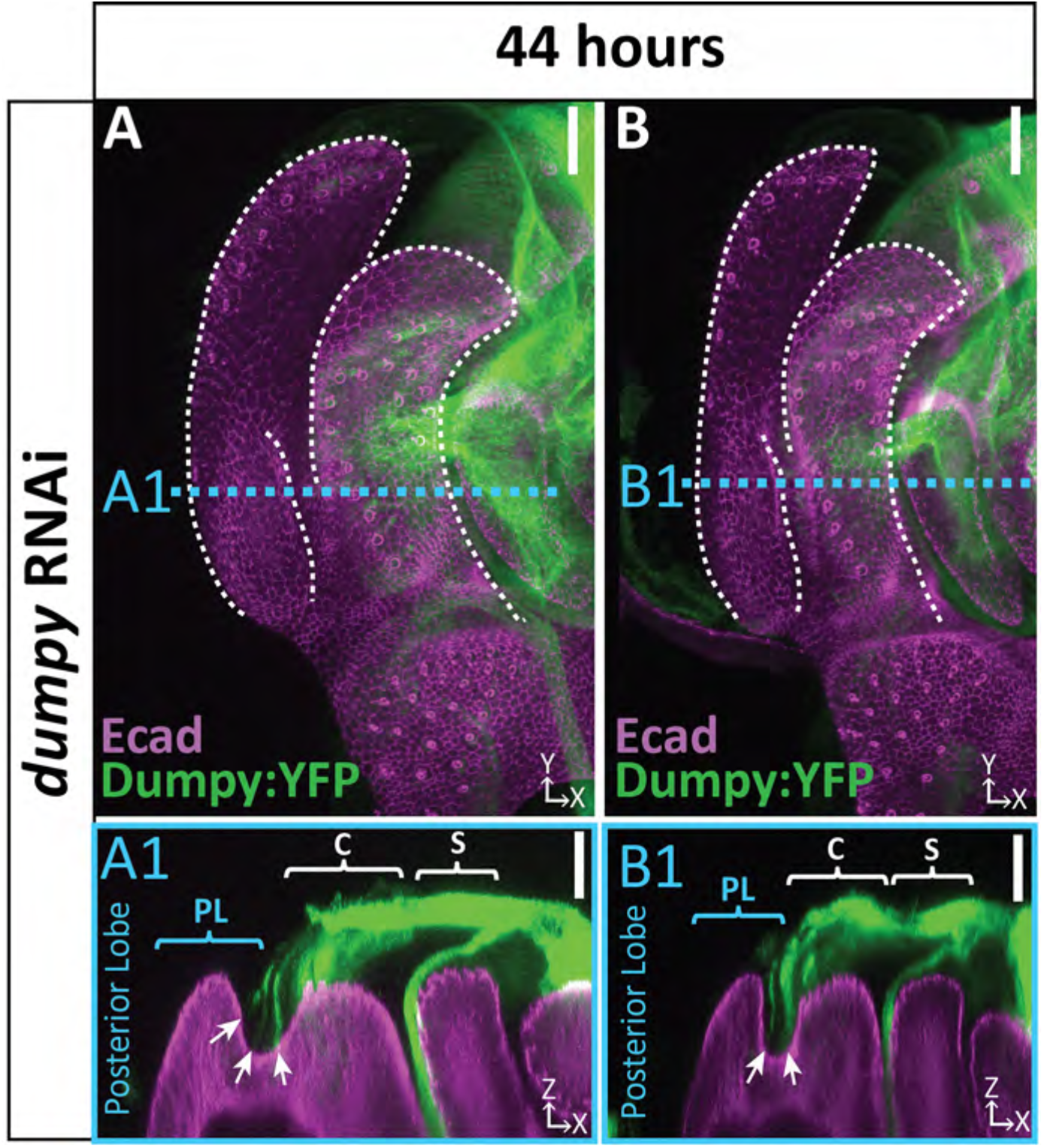
Strands of Dumpy in *dumpy* knockdown. (A & B) *dumpy* RNAi at 44 hours APF with Dumpy:YFP showing strands of Dumpy connecting to the crevice between the lateral plate and clasper (arrow). Relevant structures labeled: Lateral plate (LP) posterior lobe (PL), and clasper (C). Cross-sections are max projection of 5.434μm section to show full Dumpy connection. Scale bar, 20μm.

**Figure 8 - supplement 1.**
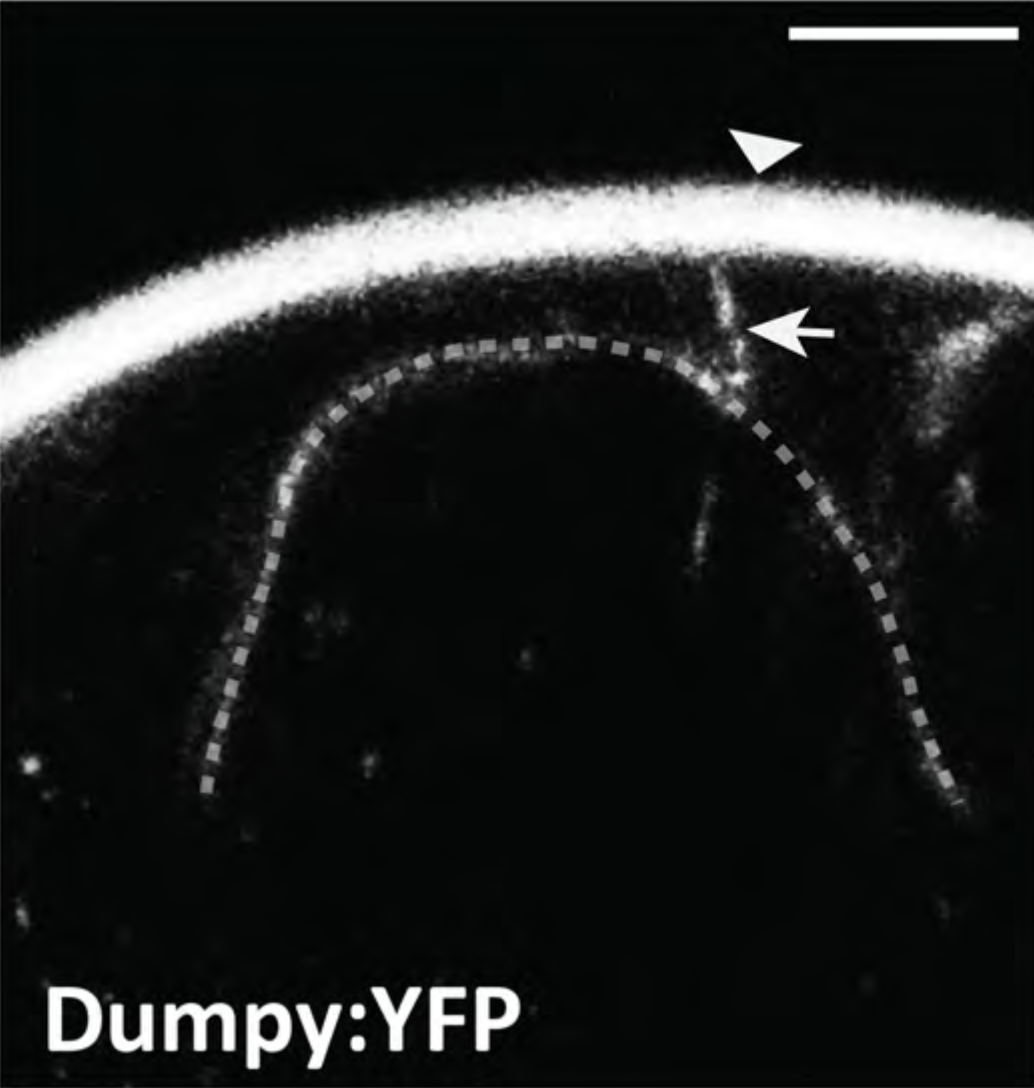
Dumpy anchors posterior spiracles to surrounding cuticle. Live imaging of Dumpy:YFP in the embryonic posterior spiracles. Posterior spiracle (dotted line) is connected to the cuticle (arrowhead) via a tether of dumpy (arrow). Scale bar, 20μm.

